# Single cell multiome profiling of pancreatic islets reveals physiological changes in cell type-specific regulation associated with diabetes risk

**DOI:** 10.1101/2024.08.03.606460

**Authors:** Hannah M Mummey, Weston Elison, Katha Korgaonkar, Ruth M Elgamal, Parul Kudtarkar, Emily Griffin, Paola Benaglio, Michael Miller, Alokkumar Jha, Jocelyn E Manning Fox, Mark I McCarthy, Sebastian Preissl, Anna L Gloyn, Patrick E MacDonald, Kyle J Gaulton

**Affiliations:** Bioinformatics and Systems Biology Program, University of California San Diego, La Jolla CA; Biomedical Sciences Program, University of California San Diego, La Jolla CA, USA; Department of Pediatrics, University of California San Diego, La Jolla CA, USA; Center for Epigenomics, University of California San Diego, La Jolla CA, USA; Department of Pediatrics, Stanford School of Medicine, Stanford University, Stanford CA, USA; Department of Pharmacology, University of Alberta, Edmonton, Alberta, Canada; Alberta Diabetes Institute, University of Alberta, Edmonton, Alberta, Canada; Wellcome Trust Center for Human Genetics, University of Oxford, Oxford, UK; Institute of Experimental and Clinical Pharmacology and Toxicology, Faculty of Medicine, University of Freiburg, Germany; Department of Genetics, Stanford School of Medicine, Stanford University, Stanford CA, USA; Stanford Diabetes Research Center, Stanford School of Medicine, Stanford, CA, USA

## Abstract

Physiological variability in pancreatic cell type gene regulation and the impact on diabetes risk is poorly understood. In this study we mapped gene regulation in pancreatic cell types using single cell multiomic (joint RNA-seq and ATAC-seq) profiling in 28 non-diabetic donors in combination with single cell data from 35 non-diabetic donors in the Human Pancreas Analysis Program. We identified widespread associations with age, sex, BMI, and HbA1c, where gene regulatory responses were highly cell type- and phenotype-specific. In beta cells, donor age associated with hypoxia, apoptosis, unfolded protein response, and external signal-dependent transcriptional regulators, while HbA1c associated with inflammatory responses and gender with chromatin organization. We identified 10.8K loci where genetic variants were QTLs for *cis* regulatory element (cRE) accessibility, including 20% with lineage- or cell type-specific effects which disrupted distinct transcription factor motifs. Type 2 diabetes and glycemic trait associated variants were enriched in both phenotype- and QTL-associated beta cell cREs, whereas type 1 diabetes showed limited enrichment. Variants at 226 diabetes and glycemic trait loci were QTLs in beta and other cell types, including 40 that were statistically colocalized, and annotating target genes of colocalized QTLs revealed genes with putatively novel roles in disease. Our findings reveal diverse responses of pancreatic cell types to phenotype and genotype in physiology, and identify pathways, networks, and genes through which physiology impacts diabetes risk.

## Introduction

Pancreatic islets are clusters of endocrine cells composed of multiple cell types that produce different hormones to maintain glucose homeostasis. A reduction in functional beta cell mass is a primary cause of type 1 and type 2 diabetes (T1D, T2D)^1–4^. The pathogenesis of T1D is characterized by immune cell-mediated death of beta cells^5^ whereas T2D is characterized by beta cell dysfunction and insulin resistance^6^. Physiological variation is a major contributor to T2D; for example, age and body mass index (BMI) are well-established risk factors for T2D^7^ and beta cells of older individuals exhibit reduced insulin secretion and expression of beta cell identity genes^8^. In addition, many genetic risk variants for T2D are also associated with measures of islet function in non-diabetic individuals such as fasting glucose, 2 hour glucose, pro-insulin levels and HOMA-B^9,10^. Physiological changes in islet cell type function have also been implicated in risk of T1D^3,4^, although the mechanisms are less well-established. A greater understanding of physiological variability within islet cell types is critical for understanding diabetes risk and heterogeneity.

The regulation of gene activity dictates cell type-specific identity and function, and many diabetes risk variants affect islet function by altering gene regulatory programs^11–13^. However, previous studies of islet gene regulation in humans had several key limitations, which prohibit deeper insight into the cell type-specific regulatory programs associated with physiological variation. Large-scale efforts to correlate genotypes to islet regulatory activity such as accessible chromatin^14^ and gene expression^15,16^ using quantitative trait locus (QTL) mapping have thus far only considered measurements in aggregate (or ‘bulk’) which obscures the effects of individual cell types. Additionally, studies that have assayed individual cell types from phenotypically diverse donors using either fluorescence-activated cell sorting (FACS)^17,18^ or single cell sequencing of single and multiomic modalities (e.g. scRNA-seq, snATAC-seq)^19–25^ have had relatively small sample sizes, particularly from non-diabetic donors, and thus limited power to detect associations. Consequently, there remains limited understanding of how phenotype and genotype impact the physiological regulation of specific cell types within islets.

Profiling islet samples from non-diabetic donors with available phenotype and genotype data using single cell genomics assays is one approach to uncover physiological variability in islet cell type-specific regulation. In this study we mapped gene regulatory programs in pancreatic islet cell types using single cell multimodal profiling (paired snRNA-seq and snATAC-seq) of 28 non-diabetic donors, which we combined with multiomic single cell profiles of up to 35 donors from the Human Pancreas Analysis Program (HPAP). We determined associations between gene regulatory programs and the phenotypes age, sex, body mass index (BMI), and glycated hemoglobin (HbA1c) levels as well as with genetic variation genome-wide. We then intersected associations with diabetes-associated variants and loci to annotate physiological mechanisms of diabetes risk.

## Results

### Single cell multimodal map of pancreatic islets in non-diabetic donors

To characterize the cell type-specific gene regulatory programs of pancreatic islets, we generated joint single nucleus RNA-seq and ATAC-seq profiles (10x Multiome) from 28 non-diabetic donors from the Alberta Diabetes Institute (ADI) IsletCore of varying age, sex, BMI, and HbA1c levels (**Supplementary Table 1**). From the same donors, we also performed microarray genotyping to characterize genome-wide genetic variation (**Figure 1a**). After applying strict quality control filters to single cell profiles (**see Methods, Supplementary Figure 1**), we performed dimension reduction and clustering of each modality individually (**Supplementary Figure 2a,b**) and then used the Weighted Nearest Neighbor (WNN) method^26^ to create a joint embedding that utilized information from both modalities. In total, the resulting multimodal map consisted of 174,819 cells (**Figure 1b**). We annotated the cell type identity of clusters using the expression level of known marker genes, which revealed endocrine beta (*INS*), alpha (*GCG*), delta (*SST*), and gamma (*PPY*) cells as well as acinar (*PRSS1*), ductal (*CFTR*), endothelial (*PLVAP*), stellate (*PDGFRB*) and immune (*PTPRC*) cells (**Figure 1c,d; Supplementary Figure 2c,d**). The accessible chromatin signal at the promoters of each marker gene further confirmed cell type assignments (**Figure 1e**). All cell types contained high quality barcodes from all donors (**Supplementary Figure 2e,f**). We provide this multimodal single cell map, along with associated resources, through interactive tools and visualizations available at http://multiome.isletgenomics.org.

**Figure 1.**
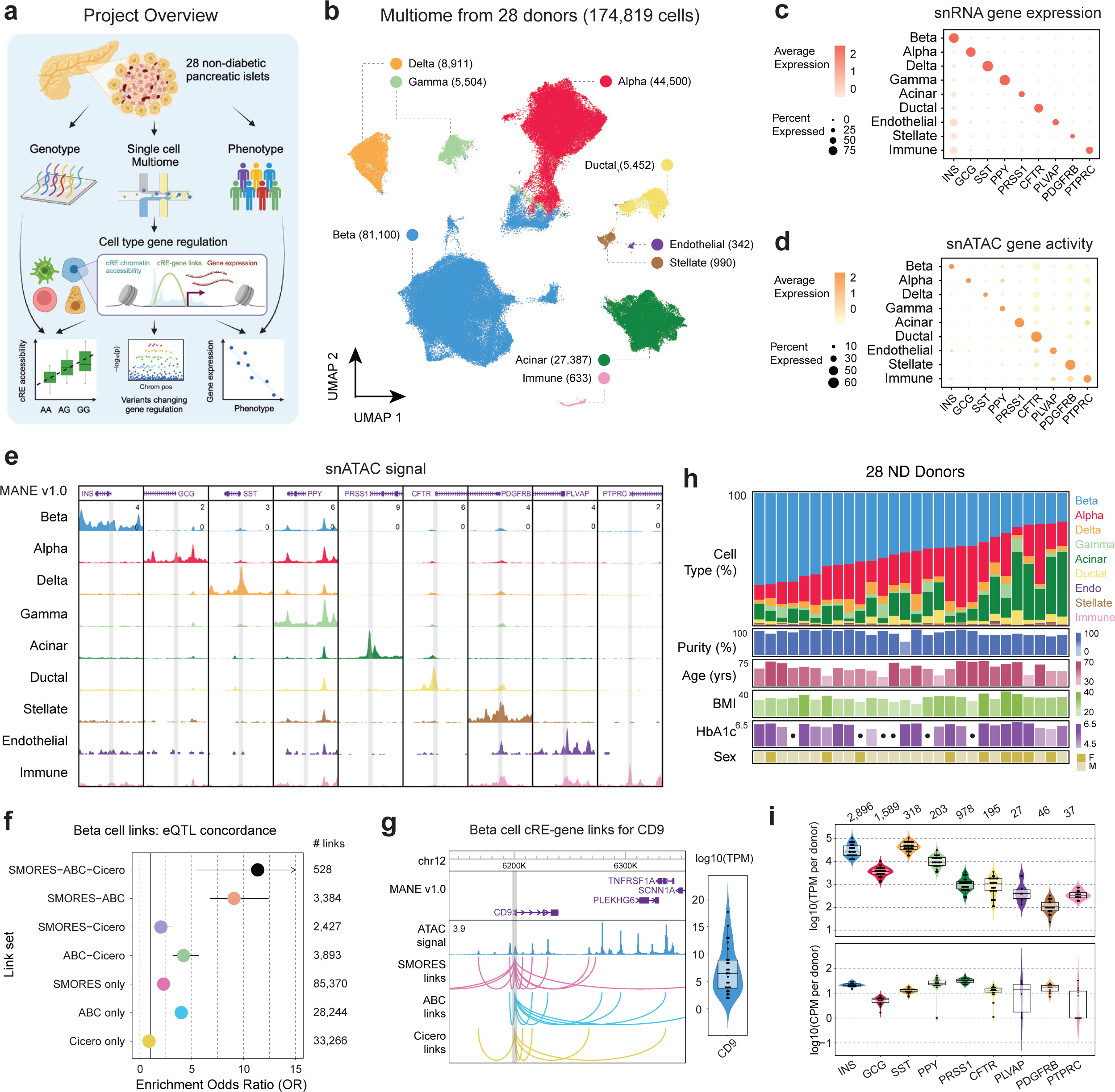
Mapping gene regulation in non-diabetic pancreatic islets with multimodal snRNA-seq and snATAC-seq. a) Project overview, 28 pancreatic islets were isolated from non-diabetic donors and genotyping and single cell Multiome was run on each. These were then used to map cell type gene regulation and how donor genotype and phenotype as well as disease risk alter regulation. Created with BioRender.com. b) UMAP projection of 174,819 barcodes post QC filtering and doublet removal. Dimensionality reduction was performed on both modalities separately and then a joint representation was created with the weighted nearest neighbors^26^ method. c) Per-cell type gene expression of islet marker genes. The dot color represents the average expression in a cell type, and the dot size represents what percent of cells in the cell type have non-zero expression. d) Per-cell type gene activity of islet marker genes. The dot color represents the average gene activity in a cell type, and the dot size represents what percent of cells in the cell type have non-zero gene activity. e) ATAC signal tracks for all cell type marker genes. Each marker gene window is a 6kb region centered on the marker gene transcription start site (TSS), and peak heights are standardized across all cell types for each region. f) Enrichment of eQTLs with the same target gene prediction in the cREs from sets of links predicted by different combinations of methods. g) Beta cell cRE-gene links between CD9 and all cREs predicted to regulate it, separated by prediction method. Right inlay shows the range of per-donor ‘pseudo’-bulk beta cell TPM values for CD9. h) Distribution of cell type proportions and phenotype information across all 28 donors, where each column represents a donor. The dots in HbA1c measurements indicate donors with missing measurements. i) Per-donor ‘pseudo’-bulk TPM and CPM values for each cell type marker gene, for the relevant cell type. Violin plots are colored based on the cell type the gene is a marker for.

We used the multimodal map to characterize gene regulatory programs of each cell type in the non-diabetic state including gene expression levels, *cis*-regulatory elements (cREs), and transcription factor (TF) binding motifs. We defined cell type gene expression and accessible chromatin levels by aggregating the profiles of individual cells for each cell type. In total, 28,862 genes were expressed (TPM>1) in any cell type with, on average, 21,165 genes expressed per cell type (range 19,715-23,857 genes per cell type) (**Supplementary Table 2**). In addition, we identified 260,281 cREs across all cell types with an average of 95,492 cREs per cell type (range 45,875-116,417 per cell type) (**Supplementary Table 3**). While we identified consistent numbers of cREs for all endocrine and exocrine cell types, several less common cell types (endothelial, immune, stellate) had <100 cells in our dataset and therefore fewer numbers of cREs due to under-sampling.

As the target genes of cRE activity are largely unknown, we next predicted cRE-target gene links in each cell type. We employed three different methods to predict cRE-target gene links for the 6 most common cell types; Activity-by-contact (ABC)^27^, Cicero^28^, and a method we developed that leverages paired single cell profiles named Single-cell MultimOdal REgulatory Scorer (SMORES) (**see Methods, Supplementary Figure 3a,b, Supplementary Data 1**). The three methods differed in both the number of linked cREs per target gene as well as the average distance between cREs and target genes (**Supplementary Figure 3c,d**). We evaluated the quality of the links from each method by testing for enrichment across three orthogonal datasets: (i) expression QTLs in pancreatic islets, (ii) 3D chromatin interactions in human pancreatic islets, and (iii) genetic fine-mapping data for T2D and fasting glucose (**Figure 1f; Supplementary Figure 3e-h**). The strongest enrichments for each dataset were for links shared among two or more methods, although there was also evidence for enrichment for links identified by ABC or SMORES only (**Figure 1f; Supplementary Figure 3e,f**). For both ABC and SMORES links these enrichments were also consistent across distance bins, even at much larger (>1Mb) distances (**Supplementary Figure 3g,h**).

In total, when considering the union of links across methods, genes had an average of 6.2 linked cREs per cell type among which 1.5 were identified by more than one method. For example, the *CD9* gene, which is a previously described marker of beta cell heterogeneity^29^, was expressed in beta cells (avg. beta cell TPM=7.02) and linked to 16 beta cell cREs across a 1Mb region including 7 links which were shared across methods (**Figure 1g**). Similarly, in acinar cells the chemokine *CXCL8* (avg. acinar cell TPM=362.26) was linked to 15 cREs including 5 which were shared across methods (**Supplementary Figure 3i**). When comparing cRE-target gene links across cell types, most links were specific to one cell type, although among the set of shared links there was a greater degree of overlap between more closely related cell types (**Supplementary Figure 3j**).

We next identified gene regulatory programs with activity specific to each cell type and which may drive cell type specialization and function. For each gene and cRE, we determined specificity in normalized profiles across cell types using Shannon entropy (**see Methods**). Genes with expression levels specific to a cell type included both known marker genes as well as, in many cases, genes that have no known function in the cell type that may be compelling targets for future studies (**Supplementary Table 4**); for example, beta cell-specific genes included *INS, G6PC2, SIX3,* and *IAPP* as well as *PLCH2, ZMAT4, WSCD2,* and *CAPN13,* and delta cell-specific genes included *SST* as well as *CADM2* and *EYS*. There were also 12,525 cREs with activity specific to a cell type, and we performed TF sequence motif enrichment using the cREs specific to each cell type with Homer (**Supplementary Table 5, Supplementary Table 6**). For TF motifs enriched in cell type-specific cREs, we correlated motif accessibility with the expression of each TF family member across samples to identify TFs likely driving cell type-specific regulation (**see Methods, Supplementary Table 6**). We identified an average of nine TFs per cell type driving cell type-specific activity (correlation FDR<.10) including *NEUROD1* and *GLIS3* in beta cells, *RFX6* in beta and delta cells, and *GATA4* in acinar cells. In gamma cells, specific cREs were enriched for TGIF family motifs and *TGIF1* expression correlated with TGIF motif accessibility, highlighting *TGIF1* as a potential novel regulator of gamma cell identity.

We next investigated the proportion of each cell type across non-diabetic donors. Endocrine cells, particularly beta and alpha cells, comprised the largest proportion of cells overall, although there was large variability in endocrine proportion across donors (**Figure 1h**). For example, the proportion of beta cells among non-diabetic donors ranged from 21.5% to 69.3%. We investigated potential technical drivers of cell type abundance, considering both biological and technical covariates (**Supplementary Figure 4a-c**). Islet isolation purity (m=.83, sd=.10), as expected, was significantly associated with abundance including increased (p=.018) beta cell proportion and decreased (p=.0074) acinar cell proportion (**Supplementary Figure 4d-f, Supplementary Table 7**). Few other technical factors strongly associated with abundance, although collagenase type was associated with ductal proportion and ischemia time was associated with alpha cell proportion (**Supplementary Figure 4g**).

We also examined variability in cell type gene expression and cRE accessibility across samples using cell type-specific ‘pseudo’-bulk profiles, focusing on marker genes for each cell type (**Figure 1i**). In beta cells, the average expression of *INS* per sample was 355,071 TPM and *INS* expression levels varied by more than an order of magnitude (sd=334,672 TPM, min=11,185 TPM, max= 844,195 TPM) across samples. We further examined the beta cell expression of a set of 16 previous reported canonical beta cell genes, and most genes were consistently expressed in all samples although several (e.g. *G6PC2, MAFA*) had wide variability in expression across samples (**Supplementary Figure 2c**). Among less common cell types (stellate, endothelial, ductal), all marker genes were expressed in every sample although with generally greater variability in the expression levels compared to more abundant cell types. We observed similar patterns in the promoter accessibility of marker genes, with the exception that some samples had complete dropout of promoter signal particularly for the rarer cell types (**Figure 1i**).

We finally examined the broad distribution of gene expression and accessible chromatin profiles of cell types across donors using principal component analysis. There was little evidence for major, systematic differences in the gene expression profiles of individual donors in any cell type (**Supplementary Figure 5**). We next investigated potential technical drivers of gene expression levels in each cell type. In line with previous studies using samples from the same biorepository^30–32^, we found a significant correlation (FDR<.10) between islet culture time and gene expression profiles, which was consistent across almost every cell type, as well as accessible chromatin profiles in many cell types (**Supplementary Table 7**).

### Physiological measures associated with islet cell type regulatory programs

To understand the implications of changes in gene regulation in pancreatic cell types on physiology, we tested for associations between donor age, BMI, HbA1c and sex and cell abundance, gene expression, cRE accessibility, and TF activity in each cell type while controlling for relevant sample and technical covariates (**see Methods**). To increase the power of these analyses we included single cell data generated in purified islets from non-diabetic donors by the Human Pancreas Analysis Program (HPAP)^33^. This included single cell RNA-seq (scRNA-seq) and single nucleus ATAC-seq (snATAC-seq) from 27 donors respectively, of which 19 donors were overlapping (35 donors in total) (**Supplementary Table 8**). For HPAP scRNA-seq we used cell type annotations from a recent publication^34^, and for HPAP snATAC-seq we performed processing, clustering, and cell type annotations for this study (**see Methods**). Donor phenotypes were evenly distributed across datasets with the exception of age, where most young donors were from HPAP whilst most older donors were from Alberta IsletCore (**Supplementary Figure 6a,b**).

We first determined whether the abundance of each cell type was associated with phenotypes. Across phenotypes including age (mean=54.42, sd=13.94), BMI (mean=28.98, sd=4.24), HbA1c (mean =5.45, sd=0.49), and biological sex (9 female, 19 male; based on self-identification and confirmed by genomic data), we identified limited evidence for significant associations (FDR<.05) with cell type abundance, although older age was significantly associated with increases in both immune cell and endothelial cell proportion (**Figure 2a, Supplementary Figure 4d**). There were also more nominal associations, including between older age and reduced acinar cell proportion and higher HbA1c level and increased ductal cell proportion (**Supplementary Table 9**).

**Figure 2.**
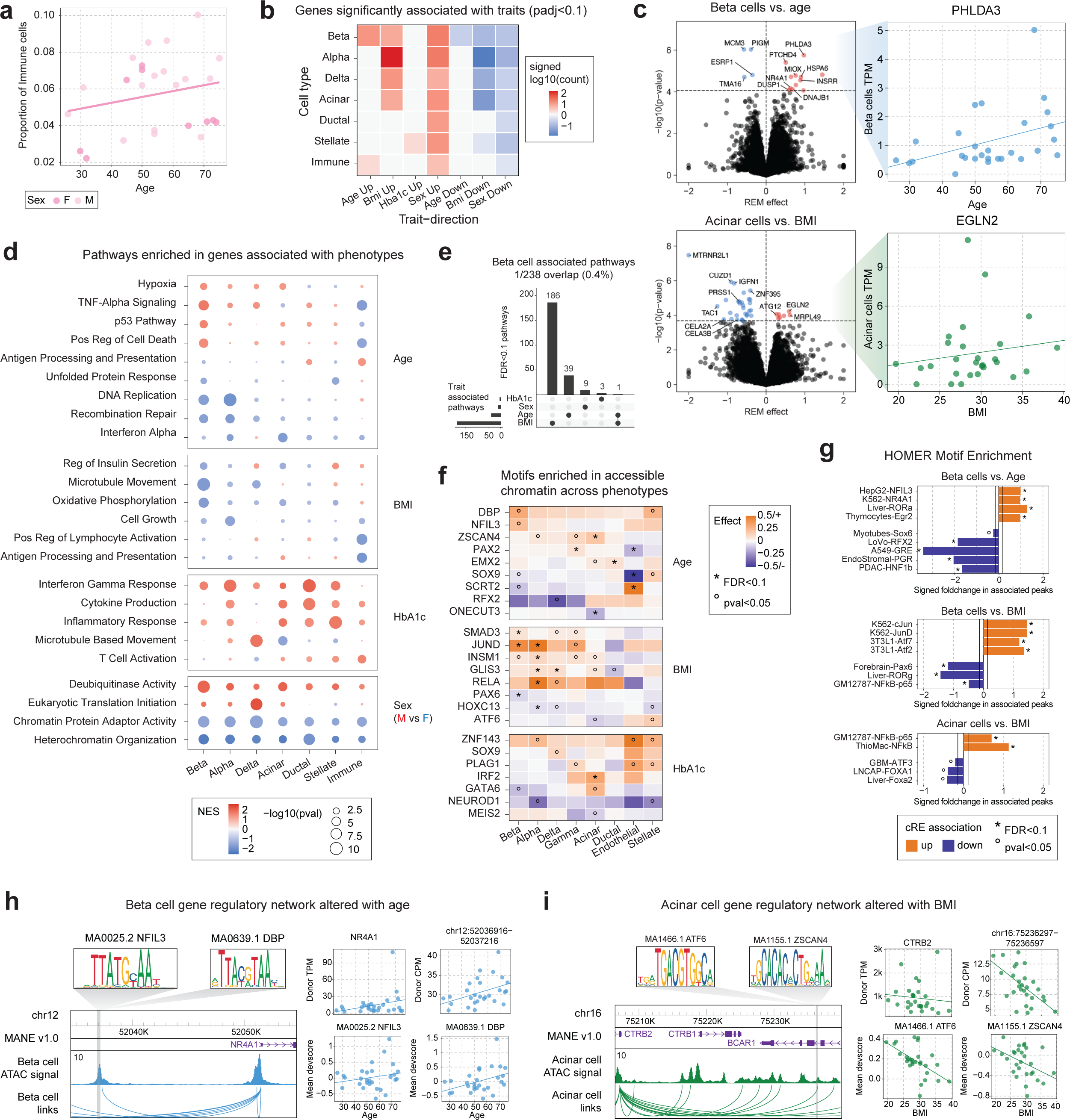
Changes in gene regulation associated with phenotype. a) Proportion of immune cells in each donor plotted against the donor age. Points are colored based on sex: light pink (M), dark pink (F). b) Summary of the number of significantly (FDR<.1) differentially expressed genes in associations between cell types and traits. Genes were divided based on the direction of differential expression and then the total number of differential genes per association was scaled with log10 and signed based on the direction. c) Left plots depict volcano plots of differential gene expression results for beta cells vs. age (top) and acinar cells vs. BMI (bottom). All genes with differential expression passing FDR<.1 are colored and the vertical line demarks the corresponding p-value cutoff. The right plots show the values used to measure associations, comparing the gene TPM values against the relevant trait, per donor in the Alberta dataset. d) Summary of pathways enriched in genes associated with phenotypes across different cell types. The color of each dot denotes the normalized enrichment score (NES) value from fGSEA and the dot size represents the -log10 of the p-value for the pathway enrichment. e) Upset plot illustrating the overlap between all pathways significantly associated with beta cells and the four phenotypes of interest. f) Heatmap of motifs nominally or significantly enriched from a meta-analysis of ChromVAR deviation scores against phenotypes. Square color reflects the effect size from meta-analysis and nominal (p<.05, °) and FDR significant (FDR<.1, *) associations are marked. g) HOMER motif enrichment results for cREs nominally (p<.01) increasing (orange) or decreasing (purple) with donor phenotypes. The solid lines denote the foldchange value associated with the top 20% of motifs tested. Nominal (p<.05, °) and FDR significant (FDR<.1, *) enrichments are marked. h-i) Example regulatory relationships between cREs and predicted target genes that show concordant changes between beta cells and age (h) and acinar cells and BMI (i). Additionally, TF motifs within the cRE with concordant changes are also shown.

We next identified gene expression levels in each cell type associated with phenotypes. In brief, we tested for association between expression and phenotypes separately for each dataset (Alberta IsletCore, HPAP) and then meta-analyzed results using a random effects model (**see Methods**). Overall, a total of 450 genes had significant association (FDR<.1) with at least one phenotype across cell types and traits (**Figure 2b, Supplementary Table 10**). In beta cells, genes associated with increased age included *PHLDA3* (FDR=.0102) (**Figure 2c, Supplementary Figure 6c**), a downstream mediator of p53^35^, heat shock genes *DNAJB1* (FDR=.094) and *HSPA6* (FDR=.035) and genes involved in stress related to reactive oxygen species *MIOX* (FDR=.035) and *DUSP1* (FDR=.094)^36^. We also identified more nominal evidence for increased expression of genes previously linked to age such as *CDKN2A* (**Supplementary Table 10**). Genes in acinar cells associated with increased BMI included *EGLN2* (FDR=.049) (**Figure 2c, Supplementary Figure 6d**), involved in hypoxia response, and *ATG12* (FDR=.057), which is involved in autophagy and apoptosis^37^. There was also significantly decreased expression of pancreatic enzymes in acinar cells in higher BMI donors such as *CELA2A* (FDR=.0813), *CELA3B* (FDR=.0813), and *PRSS1* (FDR=.024) (Supplementary Table 10).

We next performed gene set enrichment of gene associations for each trait, which revealed further insight into physiological variability in each cell type (**Figure 2d, Supplementary Table 11**). Overall, there was little overlap in the pathways significantly associated across different phenotypes within a cell type (**Figure 2e, Supplementary Figure 6e**), and slight sharing among pathways associated with a trait across cell types (**Supplementary Figure 6e-i**). In beta cells, older age was associated with an increase in hypoxia response (NES=1.55, FDR=.037), TNFa signaling (NES=1.83, FDR=1.5×10^-4^), p53 signaling (NES=1.67, FDR=.009) and cell death (NES=1.59, FDR=.031) (**Figure 2d, Supplementary Table 11**). Conversely, older age was associated with down-regulation of unfolded protein response (NES=-1.56, FDR=.086), DNA replication (NES=-1.90, FDR=.007) and recombination repair (NES=-1.75, FDR=.047) pathways (**Figure 2d, Supplementary Table 11**). Alpha cells had changes in DNA replication (NES=-2.07, FDR=1.55×10-5), cell cycle activity (NES=-1.76, FDR=.001), and recombination repair (NES=-1.86, FDR=.029) in older age, as well as decreased inflammatory signaling (interferon alpha NES=-1.79, FDR=.046) (**Figure 2d, Supplementary Table 11**). In acinar cells there were similar changes in hypoxia response (NES=1.43, FDR=.06) and inflammatory signaling in age (Interferon alpha: NES=-1.86, FDR=5.1×10-4), whereas immune cells showed marked decreases in cell death (NES=-1.65, FDR=.012) and inflammatory signaling (Interferon alpha: NES=-1.42, FDR=5.17×10-2) and increases in antigen processing and presentation (NES=1.45, FDR=.064) and cGMP binding (NES=1.45, FDR=.064), which triggers immune responses^38^.

We also observed changes in cellular pathways in the pancreas associated with BMI, HbA1c, and biological sex. Overall, increased BMI was associated with decreased activity of pathways that are key to many cell types (**Figure 2d, Supplementary Table 11**). In beta cells, increased BMI was associated with reduced expression of members of an insulin secretion pathway (NES=-1.60, FDR=1.5×10^-3^) and several related processes (microtubule movement NES=-1.89, FDR=1.11×10^-5^; calcium ion transport NES=-1.72, FDR=.053), decreased oxidative phosphorylation (NES=-1.85, FDR=2.5×10^-3^) and respiration (NES=-1.76, FDR=.0028), and nominally decreased UPR (NES=-1.46, FDR=.10). Alpha cells had decreased expression of the same processes, as well as reduced cell growth (NES=-1.66, FDR=.02). Immune cells had decreased activation (NES=-1.87, FDR=.003), interferon gamma response (NES=-1.76, FDR=.003), and antigen presentation (NES=-2.02, FDR=3.9×10^-3^). For HbA1c level and biological sex, changes in cellular function were more consistent across cell types. Higher HbA1c levels were associated with up-regulation of inflammatory signaling responses, and biological sex was associated with chromatin remodeling- and organization-related pathways, in nearly every cell type (**Figure 2d**).

We next sought to understand epigenomic changes in each cell type associated with age, sex, BMI and HbA1c. We first identified individual cREs in each cell type associated with each trait. There were 1,160 total cREs across cell types with significant association (FDR<.10) to at least one trait, most prominently for beta cell cREs and age (n=548) and BMI (n=57), alpha cell cREs and sex (n=249), and acinar cREs and HbA1c level (n=84), as well as a larger set of cREs in each cell type with more nominal (p<.01) association (**Supplementary Figure 7a,b; Supplementary Table 12**). Phenotype-associated cREs in most cell types were linked to genes with significantly concordant changes in expression for the same phenotype (**see Methods, Supplementary Figure 7c**).

To understand transcriptional regulators of epigenomic changes in phenotypes, we next estimated the accessibility of TF motifs in cells from each donor using chromVAR^39^ and tested for association between donor-level accessibility of each TF motif and phenotype (**see Methods; Supplementary Table 13**). We orthogonally performed sequence motif enrichment of cREs in each cell type with nominal association (p<.01) to each trait using Homer^40^ (**Supplementary Table 14**). In beta cells, sequence motifs for PAR-family TFs (DBP, NFIL3) had broadly increased accessibility with age and were enriched in cREs up-regulated in older age (**Figure 2f,g**). In comparison, RFX motifs had decreased accessibility, and were enriched in cREs down-regulated, in older age (**Figure 2f,g**). We also observed changes in other motifs in beta cells with age including up-regulation of NR4A1, EGR, ROR TF motifs, and down-regulation of SOX, steroid hormone receptor (GRE, PGR), HNF, and SCRT2 TF motifs, with older age (**Figure 2f,g**). The enrichment of beta cells for nuclear hormone receptors and PAR-family TFs suggests that the pathways altered in older age are driven by external signals. Other endocrine cell types showed limited enrichment of TF motifs in up-regulated cREs although RFX motifs had decreased accessibility and were enriched in cREs down-regulated with age, suggesting that reduced RFX activity is a common feature of endocrine cells in older age (**Figure 2f,g**).

Compared with age, largely distinct sets of TFs had changes in accessibility associated with BMI and HbA1c in beta cells and other cell types. For BMI, in beta cells and other endocrine cell types, JUN family TF motifs had increased accessibility and were enriched in cREs with up-regulated activity in higher BMI (**Figure 2f,g**). We also observed in endocrine and exocrine cell types increased accessibility of NFkB-related (RELA, p65) motifs in higher BMI and, in endocrine cells, increased accessibility of SMAD3, GLIS3 and INSM1 TF motifs (**Figure 2f,g**). Conversely, in beta cells, PAX6 motifs had reduced accessibility and, along with ROR, were enriched in cREs with down-regulated activity in higher BMI (**Figure 2f,g**). For HbA1c levels, there was a consistent decrease in the accessibility of NEUROD1 motifs and increase in the accessibility of several zinc fingers (ZNF143, PLAG1) in most cell types (**Figure 2f, Supplementary Figure 7g-i**). We also observed cell type-specific responses to HbA1c level, including altered IRF and MEIS2 accessibility in acinar cells (**Figure 2f**). Among cREs up-regulated in higher HbA1c levels, NFkB-related motifs were strongly enriched across cell types, in line with the increased inflammatory responses, as well as STAT motifs in beta cells (**Figure 2f, Supplementary Figure 7g-i**).

We highlighted examples of gene regulatory networks at specific loci with concordant phenotype-associated changes in activity. For example, a beta cell cRE predicted to regulate *NR4A1* had increased accessibility in older donors (log2FC=0.128, p=.0044) and contained sequence motifs for multiple PAR-family factors (**Figure 2h**). Several PAR-family TFs had nominally increased gene expression in older donors (*NFIL3*: P=.009; *DBP*: P=.006) and their motifs also had increased accessibility in older donors. Additionally, *NR4A1*, a nuclear steroid hormone receptor, had significantly higher expression (FDR=.068) and motif accessibility (FDR=1×10^-4^) in beta cells from older donors. Similarly, in acinar cells, the pancreatic enzyme *CTRB2* had nominally reduced expression in donors with high BMI (P<.026, FDR=.59), and an acinar cRE predicted to regulate *CTRB2* had decreased accessibility in higher BMI (log2FC=-0.26, p=.043) and sequence motifs for multiple TFs (ATF6, ZSCAN4) with reduced activity in high BMI (**Figure 2i**).

### Genetic variants associated with islet cell type accessible chromatin

We next determined pancreas cell type-specific gene regulatory programs affected by genetic variation in the non-diabetic state. To profile genetic variation in each sample, we performed microarray genotyping followed by genotype imputation into the TOPMed-r2 reference panel (**see Methods**). Most individuals were of European ancestry, although there were several individuals from other ancestry groups (**Supplementary Figure 8a,b**). We then identified variants genome-wide associated with cRE activity in each cell type using chromatin QTL (caQTL) mapping. In brief, for all endocrine and exocrine cell types, we tested variants in a 10kb window around each cRE for association to chromatin accessibility level across samples using RASQUAL^41^, a method that performs joint population-based (QTL) and allelic imbalance mapping. Due to the limited sample size, we were unable to perform expression QTL (eQTL) mapping in this study.

In total, we identified 10,833 cREs that had significant caQTLs (FDR<.05) in at least one cell type, with an average of 2,327 cREs per cell type (range 524-5,932) (**Figure 3a, Supplementary Figure 8c,d, Supplementary Data 2**). For comparison, we also identified 12,217 ‘bulk’ caQTLs by aggregating all ATAC-seq profiles for each sample. We performed extensive checks which confirmed the quality of the cell type caQTLs including reference allele bias, q-value distribution, lead variant proportion and cRE distance, and other metrics (**Figure 3b; Supplementary Figure 9a,b**). As expected, there was a strong correlation between the frequency of a cell type and the number of caQTLs identified for that cell type, where beta cells (n=5,923) and alpha cells (n=4,218) had the largest number of caQTLs compared to less common cell types (**SFigure 8d**). We compared caQTLs in each cell type to ‘bulk’ caQTLs in our study, as well as bulk caQTLs in pancreatic islets from a previous study^14^. There was strong concordance in the effects of cell type caQTLs with both sets of bulk caQTLs, with greater concordance for more common cell types in purified islets (beta cells: 95%, ductal cells: 82%) (**Figure 3c, Supplementary Figure 9c**).

**Figure 3.**
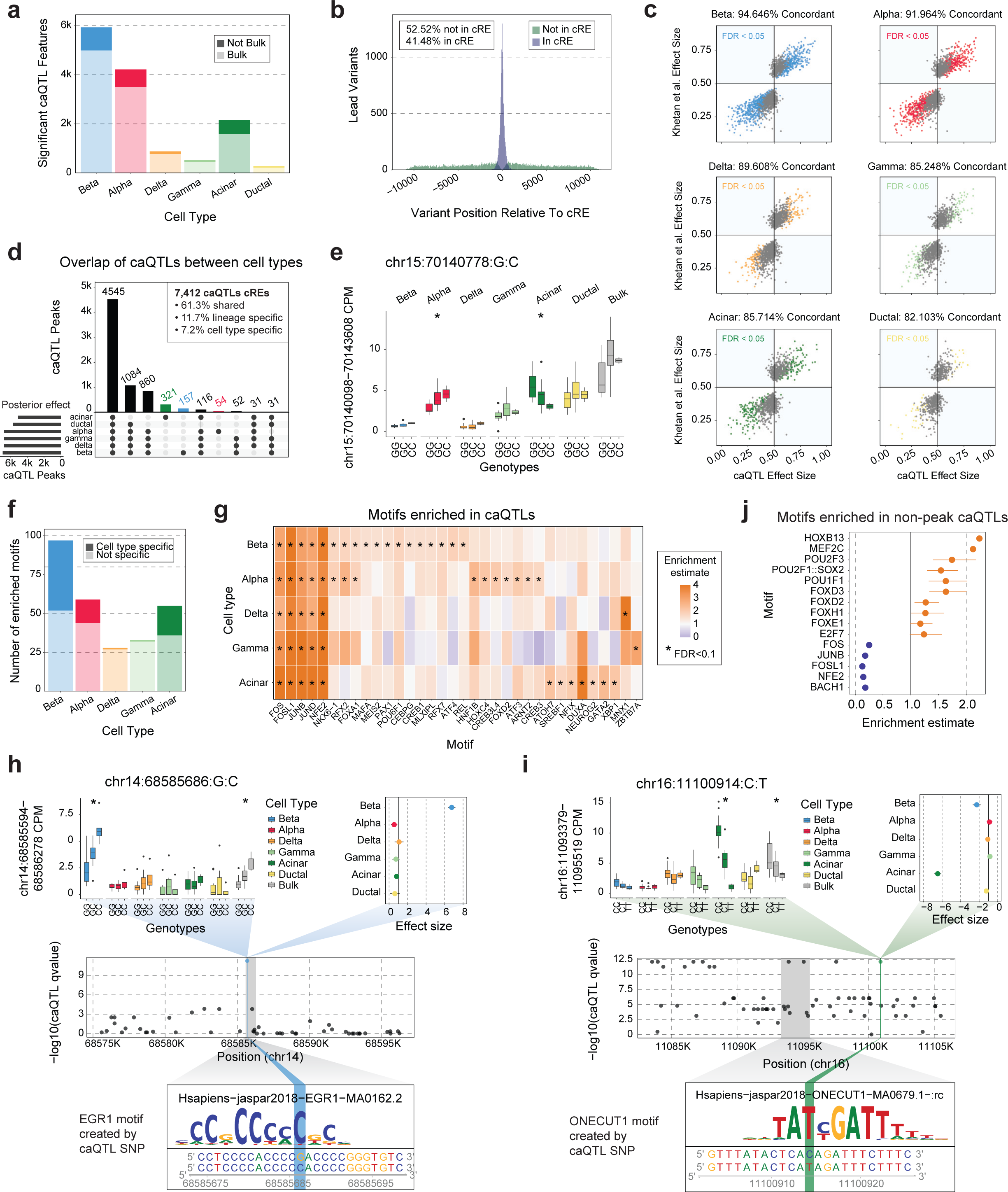
Changes in chromatin accessibility associated with genotype. a) Summary of the number of cREs with one or more significant caQTLs associated with changes in the cRE’s accessibility. These are further subdivided based on whether this cRE had significant caQTLs in our ‘pseudo’-bulk analysis (lighter color) or not (darker color). b) Distribution of the distance between a lead variant caQTL and the cRE it is associated with, colored based on whether the variant is in the cRE (blue; 41.48%) or not (green; 52.52%). c) Comparison of caQTL effect size (x-axis) and bulk islet caQTL effect sizes (y-axis) from Khetan et al^14^. Points are colored based on whether they pass FDR significance in our results, and the percentage of variants with concordant directional effects is listed at the top of each plot. d) Upset plot showing the overlap between peaks with significant caQTLs in different cell types based on mashR’s posterior effect calculation. The top 10 intersection sets are displayed. e) Example of a caQTL (chr15:70140778:G:C) that was significant in both alpha and acinar cells but had different directions of effect between the two cell types. Colored-coded boxplots show association in different cell types and white boxplots show our ‘pseudo’-bulk islets. Significant caQTLs (FDR<.05) are indicated with *. f) Summary of the number of enriched motifs in caQTLs for each cell type, color shade indicates whether this motif was only enriched in caQTLs for that cell type (dark) or was enriched in multiple cell types (light). g) Heatmap of the enrichment or depletion of motifs in caQTLs by cell type. Significant enrichments (FDR<.1) are marked with *. h-i) Cell type specific caQTLs for beta cells (h: chr14:68585686:G:C) and acinar cells (i: chr16:11100914:C:T). Top left: colored-coded boxplots show association in different cell types and white boxplots show our ‘pseudo’-bulk islets. Significant caQTLs (FDR<.05) are indicated with *. Top right: mashR cell type posterior effect sizes for the caQTL in each major cell type. Middle: All caQTL p-values for the cRE of interest (coordinates marked in grey box). The caQTL of interest is colored blue (h) or green (i). Bottom: TF motif created by caQTL alternative allele and enriched in caQTL by motifbreakR. j) Enrichment and depletion of motifs in caQTL lead SNPs that do not fall into a known cRE.

We next determined the extent of sharing of caQTLs across different pancreatic cell types. We used multivariate adaptive shrinking (MASH)^42^ to re-estimate the effects and significance of caQTL variants by leveraging information across cell types, and then categorized caQTLs based on their effects (FDR<.05) across cell types (**see Methods, Supplementary Data 3**). The majority of caQTLs were shared between more than one cell type (92.8%), with a greater degree of sharing among more closely related cell types, and 61.3% of caQTLs were shared across all six tested cell types (**Figure 3d**). Among the remaining caQTLs, 11.7% had lineage-specific and 7.2% had cell type-specific effects (**Figure 3d**). Within shared caQTLs, a small proportion (1.16%) had opposed effects in different cell types. For example, variant rs56350203 (15-70140778-G-C) was a caQTL for a cRE upstream of the *TLE3* gene where the G allele had increased accessibility in acinar cells (effect=.70, FDR=9.7×10^-13^) but decreased activity in alpha cells (effect=.37, FDR=1.9×10^-12^) (**Figure 3e**), and no corresponding association in ‘bulk’ data (effect=.53, FDR>.05). The alleles of this variant were predicted to bind motifs for different TFs (G allele-RBPJ, C allele-GATA) enriched in acinar and alpha cells, respectively.

We investigated further the subset of caQTLs with cell type-specific effects on cRE activity. Overall, most caQTLs specific to a cell type were for cREs active in additional cell types, which was consistent across different cell types (acinar: 71.3%, beta: 89.8%, alpha: 90.7%). We speculated that cell type-specific caQTLs are often due to variants altering binding of TFs with cell type-restricted expression or activity. We therefore next identified TF sequence motifs preferentially disrupted by caQTL variants in each cell type using motifbreakR^43^ (**see Methods, Figure 3f**). The strongest motif enrichments in each cell type were for FOS and JUN family TF motifs, which likely represent shared effects across cell types, and we also identified TF motifs with linage-specific enrichment, such as RFX and FOXA motifs in endocrine cells, and cell type-specific enrichment (**Figure 3g, Supplementary Table 15**). Variants with cell type-specific effects included rs2253317 (14-68585686-G-C) which had beta cell-specific effects on a cRE intronic to the *RAD51B* gene and rs12927355 (16-11100914-C-T) which had acinar-specific effects on a cRE intronic to the *CLEC16A* gene, both of which had allele-specific binding to TFs (**Figure 3h,i**).

Most cell type caQTLs (86.79%) had at least one significantly associated variant directly overlapping the cRE, which was consistent across caQTLs in each cell type (associated: 85.4-88.8%), and thus many of these caQTLs likely impact cRE activity by altering binding of TFs within the cRE. At about half of the remaining caQTLs (6.2%) at least one significantly associated variant lies within a different cRE. For the remaining subset of caQTLs without associated variants in cRE boundaries, we determined whether they may affect distinct aspects of genomic regulation. We identified sequence motifs enriched in non-cRE caQTLs in each cell type compared to cRE-overlapping caQTLs. Sequence motifs significantly enriched in non-cRE caQTLs included many TFs with known pioneer activity such as FOX family TFs, POU family TFs, and HOXB13 while, conversely, non-cRE caQTLs were relatively depleted in FOS and JUN family TF motifs (**Figure 3j**, **Supplementary Table 15**). This highlights potentially distinct classes of variants affecting cRE activity for variants outside of cRE boundaries.

### Physiological changes in islet cell type regulation associated with diabetes risk

Finally, we examined how phenotypic and genetic variability in pancreatic cell type gene regulation in the non-diabetic state impacts risk of T1D and T2D. We first determined whether cell type cREs associated with each phenotype were enriched for variants associated with T1D, T2D and glycemic traits (**see Methods**). Beta cell cREs associated with HbA1c were significantly enriched for T2D risk variants (fold-enrich=4.26, FDR=.024), and beta cell cREs associated with BMI showed more nominal enrichment (fold-enrich=2.9, p=.017) (**Figure 4a, Supplementary Table 16**). Fine-mapped variants at 29 T2D signals overlap a phenotype-associated beta cell cRE (**Supplementary Figure 10a**). For example, causal T2D variants at the *TCF7L2* (rs7903146, PIP=.99) and *CDK2NA/B* (rs10757282, PIP=.99) loci map in beta cell cREs with reduced accessibility in higher BMI (**Figure 4b**). Outside of beta cells, there were few enrichments for T2D except for one nominal one with cREs associated with age in gamma cells (fold-enrich=4.2, p=.03) (**Figure 4a**). There was also limited enrichment for T1D associated variants, except for ductal cell cREs with age (fold-enrich=7.09, p=.041) and delta cell cREs with sex (fold-enrich=9.96, p=.009) (**Supplementary Figure 10b**). For glycemic traits, beta cell cREs associated with sex were nominally enriched for HbA1c associated variants (fold-enrich=5.23, p=.04) and beta cell cREs associated with BMI were enriched for fasting glucose (fold-enrich=6.09, p=.007) and insulin (fold-enrich=6.02, p=.049) level variants (**Supplementary Figure 10c-f**).

**Figure 4.**
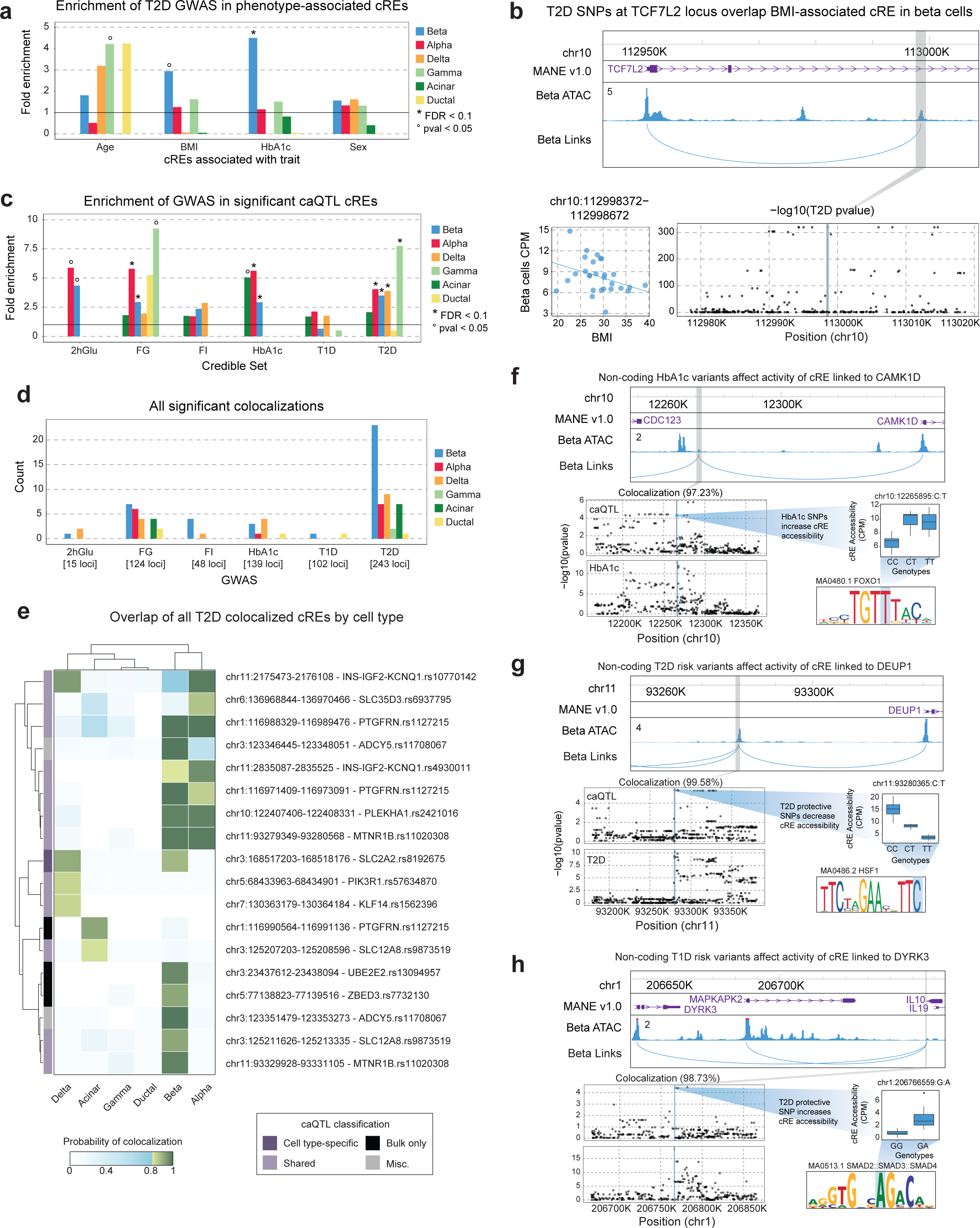
Enrichment of disease risk in changes in gene regulation associated with phenotype and genotype. a) Enrichment of SNPs in the T2D credible set that overlap cREs associated with age, BMI, HbA1c and sex, in the six largest cell types. Nominally significant enrichments (pval<.05) are indicated with °. b) Example of a BMI-associated cRE in beta cells which contains T2D risk variants at the TCF7L2 locus. Top: locus plot showing the cRE chr10:112998372-112998672 that contains T2D risk SNPs, and it’s predicted target gene TCF7L2. Bottom left: association between donor BMI and the accessibility of the cRE in beta cells per donor. Bottom right: T2D summary statistics in the TCF7L2 locus. The grey box indicates the cRE, and the blue dot and line indicates the SNP that overlaps the cRE. c) Enrichment of SNPs in the glycemic trait, T1D and T2D credible set that overlap cREs that have a significant caQTL in each cell type. FDR significant enrichments (FDR<.1) are indicated with * and nominally significant enrichments (pval<.05) are indicated with °. d) Summary of the number of statistically significant (PIP>0.8) colocalizations between cell type caQTLs and summary statistics for glycemic traits, T1D, and T2D. e) The PIP for each T2D colocalized cRE across all cell types. Non-significant colocalizations are colored in blue and significant colocalizations are colored in green. Row names indicate the nearest credible set to the cRE that overlaps T2D risk. The vertical color bar indicates the cell type classifications of caQTLs based on mashR results, with Misc referring to anything not covered by the other categories. f-h) Example colocalizations between T2D, T1D, or glycemic trait association and beta cell caQTLs predicted to regulate (f) CAMK1D, (g) DEUP1, and (h) DYRK3. Top: locus plot showing the cRE with colocalized caQTLs (grey box), and predicted target gene links. Bottom left: caQTL and GWAS summary statistics for the region around the cRE. Middle right: box plots for a representative SNP that is a caQTL for the cRE and directly overlaps a GWAS variant. Bottom right: Motif created by the example caQTL with the caQTL created base highlighted in blue.

We then determined whether cell type cREs associated with genotype (caQTLs) were enriched for diabetes and glycemic trait associated variants. Endocrine cell type caQTL cREs were all significantly enriched for T2D risk variants (beta fold-enrich=3.49, FDR<.0001; alpha fold-enrich=4.04, FDR<.0001; delta fold-enrich=3.9, FDR=.097; gamma fold-enrich=7.7, FDR=.012) (**Figure 4c, Supplementary Table 17**). There was no corresponding enrichment of T2D risk variants in exocrine cell type caQTLs, nor was there enrichment of T1D risk variants in caQTLs for any cell type (p<.05) (**Figure 4c**). When considering caQTLs with cell type-specific effects, only beta cell-specific caQTLs showed nominal enrichment for T2D risk variants (fold-enrich=2.6, p=.13) (**Supplementary Figure 10g,h**). For T1D, there were again no significant enrichments in cell type-specific caQTLs (**Supplementary Figure 10h**). The caQTLs with shared effects across cell types were significantly enriched for variants associated with most traits including fasting glucose (fold-enrich=6.5, FDR=.0085), HbA1c (fold-enrich=6.0, FDR=.011), and T2D (fold-enrich=6.8, FDR<.0001) (**Supplementary Figure 10h**).

We next investigated specific diabetes- and glycemia-associated loci that alter regulation of pancreatic cell types in physiology. Across all known T1D, T2D and fasting glycemia loci, candidate causal variants at 226 total loci (87 T2D, 54 T1D, 42 fasting glucose, 13 fasting insulin, 5 2-hour glucose, 34 HbA1c) were cell type caQTLs (**Supplementary Table 18**). We further identified diabetes and glycemic trait association signals with evidence for a shared causal variant with cell type caQTLs using statistical colocalization (**see Methods**). Across all cell types, there were 40 loci with a shared causal variant underlying both trait and caQTL association (posterior probability [PP]>.80) (**Figure 4d; Supplementary Figure 11a,b; Supplementary Table 19**). We annotated the putative gene targets of colocalized trait and caQTL signals using cRE-target gene links and at 26 colocalized caQTLs there was at least one predicted target gene of the cRE (**Supplementary Table 19**).

The largest number of shared variants were between T2D and glycemic trait association signals and beta cell caQTLs (**Figure 4e, Supplementary Figure 11c,d**), including multiple loci with previously reported effects on beta cell regulation such as *CDC123/CAMK1D, ADCY5,* and *ZBED3/PDE8B*^44–46^. At the *CDC123/CAMK1D* locus, HbA1c level-associated variants were associated with increased activity of a beta cell cRE predicted to regulate *CAMK1D* expression, and these same variants are also associated with T2D risk (**Figure 4f**). At other T2D loci, colocalized caQTLs provide new hypotheses about beta cell regulatory programs and genes involved in T2D. For example, risk alleles at a T2D signal downstream of *MTNR1B* were associated with increased activity of a beta cell cRE upstream of *DEUP1* predicted to regulate *DEUP1* expression (**Figure 4g**). In other examples, at the *PTGFRN* locus T2D risk alleles were associated with decreased activity of a beta cell cRE predicted to regulate *CD101* expression, among other genes, and at the *UBE2E2* locus T2D risk alleles were associated with increased activity of a beta cell cRE predicted to regulate *UBE2E2* expression (**Supplementary Table 19**). Furthermore, T2D-associated variants showed evidence for association with the expression of predicted target genes (*DEUP1* P=4.3×10^-85^; *CD101* P=1.9×10^-14^; *UBE2E2* P=1.5×10^-3^) in bulk islet eQTL data^16^, supporting these genes as potential targets of T2D-associated cRE activity.

Outside of beta cells, several caQTLs colocalized with T2D and glycemic trait association signals in other cell types and which highlight potential mechanisms of T2D risk in these cell types. Four T2D signals colocalized with non-beta endocrine caQTLs that were not colocalized in beta cells (**Figure 4e**). We determined whether these colocalized caQTLs had weaker evidence for colocalization in beta cells, and only one signal at *INS/IGF2/KCNQ1* was just below the colocalization threshold (beta cell PP=.78) (**Figure 4e**). We also identified largely distinct sets of caQTLs colocalized with T2D association signals in exocrine cells compared to beta or other endocrine cell types. As most caQTLs had shared effects across cell types, caQTLs colocalized only in specific cell types in some cases likely reflect differences in association strength, and thus colocalization power, across cell types (**Figure 4e, Supplementary Figure 11e-f**). In addition, we identified colocalizations with endocrine and exocrine cell type caQTLs at loci associated with glycemic traits, for example at the *GRB10* and *VEGFA* loci (**Supplementary Figure 11c,d**).

Although T1D risk variants were not broadly enriched in pancreas cell type caQTLs, we did identify several shared signals between T1D risk and pancreas cell type caQTLs, highlighting physiological processes in the pancreas that may affect T1D risk at specific loci. In beta cells, caQTLs were colocalized with T1D association at the *IL10* locus (**Figure 4h**) as well as, just below the shared probability threshold, the *MAPT* locus (PP=.78). At the *IL10* locus, the beta cell cRE affected by T1D risk variants was predicted to regulate *DYRK3*, which is involved in stress granule formation and mTOR signaling and inhibition of DYRK family members induces beta cell proliferation^47^ (**Figure 4h**). We also identified several caQTLs colocalized with T1D associated variants in other cell types, such as ductal cells. In addition, although not formally colocalized, significant caQTLs for candidate variants at other T1D loci highlighted additional physiological processes potentially involved in T1D risk; for example, candidate variants were beta cell caQTLs at the *INS* and *DLK1* loci and acinar cell caQTLs at the *CTRB1/2* locus.

## Discussion

Single cell multiomic profiling of the pancreas in physiology revealed substantial diversity in transcriptional responses across different cell types and phenotypes. Age, obesity and HbA1c are all major risk factors for T2D, and our findings provide insight into how these phenotypes drive T2D risk by altering islet cell type regulation. Beta cells in older age donors had increased expression of cell death-related genes, as well as hypoxia and inflammatory response and senescence-related genes which all contribute to beta cell dysfunction and death^48–59^. In addition, beta cells have reduced UPR with age, which acts downstream of ER stress and is linked to beta cell de-differentiation and dysfunction^60–62^. A recent study identified increased ER stress, autophagy and transcription in beta cells in older age^8^. Although we found limited evidence for increased ER stress response directly in our results, this may also reflect differences between cellular and nuclear expression profiles. The effects of obesity on T2D risk are largely mediated through insulin resistance in peripheral tissues, which in turn can increase demands for insulin secretion from beta cells^63–65^. In this study beta cells in higher BMI had reduced expression of genes involved in insulin secretion and related pathways, which may reflect effects of elevated glucose and lipid levels on beta cells directly^66–68^. Finally, in higher HbA1c, multiple cell types including beta cells had marked up-regulation of interferon signaling and inflammation, in line with previous reports^69^.

Our study also revealed distinct epigenomic drivers of transcriptional activity in physiology across cell types and phenotypes, many of which act downstream of external stimuli. In beta cells, older age was associated with increased activity of TFs that mediate cellular responses to external signals and stimuli. Several PAR bZIP family TFs (NFIL3, DBP, HLF) had increased age-related activity in beta cells, and these TFs are regulated by circadian rhythms^70^ which are dysregulated in older age^71,72^ and have been linked to impaired beta cell function, survival and oxidative stress response^63–65^. Supporting the role of altered PAR TF activity in T2D, a gene network regulated by DBP in beta cells is increased in pre-T2D and T2D donors^25^. Age also altered the activity of nuclear receptors such as NR4A1, NR1F1 (RORα) and the steroid hormone receptors GR and PGR. Age-related changes in beta cells therefore appear to be largely in response to external stimuli, in addition to intrinsic factors, and these signals could represent areas for therapeutic modulation to enhance beta cell survival and function. In higher BMI, beta cells had increased SMAD3 activity, which is linked to reduced insulin secretion and beta cell function^73^, as well as JUN activity, which is linked to metabolic stress response^74^. Interestingly, we also found that distinct sets of core beta cell TFs had decreased activity across different phenotypes, including RFX and HNF1 in age, PAX in BMI and NEUROD1 in HbA1c. There were also strong epigenomic responses of other cell types to phenotypes; for example, increased IRF activity in acinar cells in higher HbA1c in line with the increases in interferon signaling.

Previous studies of QTLs in pancreatic islets have relied on bulk measurements^14^, and our study provides the first insight into genetic effects on gene regulation in different islet cell types. The majority of caQTLs were shared across cell types or lineages, including risk variants at many diabetes risk loci, which may complicate understanding the specific cell types through which risk variants affect disease. Nearly 20% of chromatin QTLs had either lineage-specific or cell type-specific effects, and the specificity of QTL activity in many cases appears due to altered binding of transcription factors with cell type-restricted activity. We also identified transcriptional regulators affected by QTLs that map outside of cRE boundaries, and these variants preferentially alter binding of pioneer TFs in flanking regions of closed chromatin which may cause changes in the accessibility of nearby elements. One key limitation of our study is the limited sample size for QTL mapping, and larger samples will be necessary to map gene eQTLs in each cell type, identify independent signals for both caQTLs and eQTLs, and improve power for colocalization analyses. In addition, QTL mapping studies performed in more diverse contexts, such as in response to environmental stimuli and across developmental stages^75,76^, will reveal additional classes of QTLs in islet cell types.

Chromatin QTL mapping revealed T2D loci that affect beta cell regulation as well as their direction of effect. At several loci with established roles in beta cell function such as *TCF7L2* and *CDKN2A/B*^77–79^, cREs affected by T2D risk variants were also associated with donor phenotypes, and therefore risk variant activity may interact with disease-relevant phenotypic changes. We also identified putative novel mechanisms of T2D risk in beta cells. At a T2D signal on chromosome 11, risk variants increase the activity of a beta cell cRE linked to *DEUP1* as well as *DEUP1* expression in islets. *DEUP1* is a component of cell division machinery^80^ and, given *in vitro* evidence that DEUP1 can self-assemble^81^, increased *DEUP1* activity and assembly may alter signal reception or microtubule dynamics^82^. Other genes linked to beta cell cREs affected by T2D risk variants include *UBE2E2,* which alters insulin secretion in mouse beta cells^83^, and *CD101,* which has no established role in beta cells. In addition to target genes, the specific cREs affecting disease risk may themselves represent areas for therapeutic developing using epigenome editing techniques such as CRISPR interference. While we identified the largest number of shared caQTLs in beta cells, colocalization of T2D signals with caQTLs in other endocrine and exocrine cell types highlight potential roles for these cell types in T2D.

In comparison to T2D, we found limited evidence that physiological changes in beta cell regulation have a major impact on risk of T1D. We profiled a relatively limited number of phenotypes in this study and examining associations in an expanded set of phenotypes, particularly from juvenile donors, may help uncover relationships between physiology and T1D risk. In addition, risk of T1D in beta cells and other islet cell types may more often depend on responses to specific environmental triggers, such as proinflammatory cytokines and viral infection^75,84–86^. Despite the lack of overall enrichment for T1D, we did find some evidence for mechanisms mediating T1D risk in the pancreas at specific loci. For example, T1D associated variants at the *IL10* locus affected beta cell regulation and were linked to *DYRK3*, which is involved in the formation of stress granules that contain repressed mRNA^87^. Beta cell release of stress granules may affect T1D risk by triggering immune cells and enhancing autoimmune responses^88^. In addition, T1D associated variants at the *INS* locus affected activity of a cRE upstream of *INS* and thus may impact the regulation of insulin and other genes. Notably, the risk alleles for T1D at this locus increase cRE activity, which may support a role for insulin hypersecretion in T1D risk^89^.

As most diabetes risk variants affect the activity of cREs distal to gene promoters, defining the gene targets of cRE activity remains a major barrier to interpreting the mechanisms of disease risk loci. In our study, we utilized multiple methods to predict cRE-target gene links including ABC^27^, Cicero^28^, and a novel method we developed, SMORES, that leverages single cell multimodal profiles. Links supported by multiple different methods showed the strongest enrichments in orthogonal data such as expression QTLs, 3D chromatin conformation and GWAS fine-mapping data, most prominently for links shared between ABC and SMORES. Future studies may therefore benefit from using an ensemble of multiple methods to define relationships between cREs and target genes. These efforts will be facilitated through more systemic validation of cRE-target gene links in pancreatic cell types, for example by using CRISPR-based screens^27^ or heritability^90^, to help define the specificity and sensitivity of each linking methods. Futher studies are then needed to link changes in gene expression to disease-relevant cellular function.

Overall, mapping phenotypic and genotypic variability in gene regulatory programs in islet and other pancreatic cell types in physiology provided key insight into how islet physiology affects risk of developing diabetes. More broadly, our study provides a framework for how single cell multiomic profiling can be used to describe normal variability in cell type-specific gene regulation and its role in individual risk of complex disease.

## Methods

### Islet sample collection and processing

Human islets isolated according to established protocols (dx.doi.org/10.17504/protocols.io.bt55nq86) at the Alberta Diabetes Institute IsletCore and banked in aliquots of ∼2,000 islet equivalents by snap freezing. Islets from 28 non-diabetic donors were used in the present study (**Supplementary Table 1**). For all donors we obtained phenotypic information on their age, body mass index (BMI) and sex. For twenty-three of the donors, we also obtained measurements of glycated hemoglobin levels from the hemoglobin A1c (HbA1c) test. The processing and banking of islets for research was approved by the Human Research Ethics Board at the University of Alberta (Pro00013094) and all donors’ families gave written informed consent for the use of tissue in research. Islet studies in San Diego were approved and considered exempt by the Institutional Review Board of the University of California San Diego.

### Generation & analysis of genotype data

All samples were genotyped using the Illumina InfiniumOmni2-5Exome array, which measures approximately 2.6 million single nucleotide variants (SNV) sites. Raw genotyping data underwent quality control, including filtering out SNVs with excess missing genotypes (>2%) and samples with discordant genetic vs. recorded sex. The filtered data were then imputed using the TOPMed Imputation Server (r2 panel). Following quality control and imputation, 292 million variants, including 15,517,491 high-quality variants with imputation score (R2) > 0.3, were available for downstream analysis. The workflow included several steps: (i) base calling with initial raw data processing using BED, BIM, and FAM files, (ii) sex chromosome consistency checks using Plink and BCFtools, (iii) conversion to VCF format, (iv) chromosome-specific splitting and (iv) uploading the split data to the TOPMED imputation server, followed by downloading the imputed VCF files for chromosomes 1-22.

### Single cell multiome assays

Islets were thawed and resuspended in 1 mL wash buffer (10 mM Tris-HCL (pH 7.4), 10 mM NaCl, 3 mM MgCl2, 0.1% Tween-20 (Sigma), 1% fatty acid-free BSA (Proliant, 68700), 1 mM DTT (Sigma), 1x protease inhibitors (Thermo Fisher Scientific, PIA32965), 1 U/µl RNAsin (Promega, N2515) in molecular biology-grade water) and dounce homogenized with a tight-fitting pestle (Wheaton, EF24835AA) for 10-20 strokes. Homogenized islets were filtered (30 µm filter, CellTrics, Sysmex) and pelleted with a swinging bucket centrifuge (500 x g, 5 min, 4°C; 5920R, Eppendorf). Nuclei were resuspended in 400 µL of sorting buffer (1% FA-free BSA, 1x protease inhibitors (Thermo Fisher Scientific, PIA32965), 1 U/µl RNAsin (Promega, N2515) in PBS) and stained with 7-AAD (1 µM; Thermo Fisher Scientific, A1310).

In total 120k nuclei were sorted using an SH800 sorter (Sony) into 87.5μl of collection buffer (1 U/µl RNAsin (Promega, N2515), 5% fatty acid-free BSA (Proliant, 68700) in PBS). The nuclei suspension was mixed in a ratio of 4:1 with 5x permeabilization buffer (50 mM Tris-HCL (pH 7.4), 50 mM NaCl, 15 mM MgCl2, 0.5% Tween-20 (Sigma), 0.5% IGEPAL-CA630 (Sigma), 0.05% Digitonin (Promega), 5% fatty acid-free BSA (Proliant, 68700), 5 mM DTT (Sigma), 5x protease inhibitors (Thermo Fisher Scientific, PIA32965), 1 U/µl RNAsin (Promega, N2515) in molecular biology-grade water) and incubated on ice for 1 minute before pelleting with a swinging-bucket centrifuge (500 x g, 5 min, 4°C; 5920R, Eppendorf). The supernatant was gently removed and ∼50 µl were left behind to increase nuclei recovery. Next, 650 µl of wash buffer (10 mM Tris-HCL (pH 7.4), 10 mM NaCl, 3 mM MgCl2, 0.1% Tween-20 (Sigma), 1% fatty acid-free BSA (Proliant, 68700), 1 mM DTT (Sigma), 1x protease inhibitors (Thermo Fisher Scientific, PIA32965), 1 U/µl RNAsin (Promega, N2515) in molecular biology-grade water) was added without disturbing the pellet and nuclei were pelleted again with a swinging bucket centrifuge (500 x g, 5 min, 4°C; 5920R, Eppendorf). The supernatant was removed without disturbing the pellet and 7-10 µl of 1x Nuclei Buffer (10x Genomics) was added and nuclei were resuspended. Nuclei were counted using a hemocytometer, and approximately 18k nuclei were used as input for tagmentation.

Single-cell Multiome libraries were generated following manufacturer instructions (Chromium Next GEM Single-cell Multiome ATAC + Gene Expression Reagent Bundle, 1000283; Chromium Next GEM Chip J Single cell, 1000234; Dual Index Kit TT Set A, 1000215; Single Index Kit N Set A, 1000212; 10x Genomics) with 7 cycles for ATAC index PCR, 7 cycles for cDNA amplification, 13-16 cycles for RNA index PCR. Final libraries were quantified using a Qubit fluorimeter (Life Technologies) and the size distribution was checked using Tapestation (High Sensitivity D1000, Agilent). Libraries were sequenced on NextSeq 500 and NovaSeq 6000 (Illumina) with read lengths (Read1+Index1+Index2+Read2): ATAC (NovaSeq 6000) 50+8+24+50; ATAC (NextSeq 500 with custom recipe) 50+8+16+50; RNA (NextSeq 500, NovaSeq 6000): 28+10+10+90.

### Data quantification and alignment

10x Genomics Cell Ranger ARC (v2.0.0) was used to pre-process all raw sequence data. The cellranger count command was used to align the sequencing reads to the genome reference GRCh38 with the reference annotation GENCODE v32. Bam files and feature barcode count matrices were generated.

### Processing single nucleus per-donor counts data

Before merging data from all donors, we individually processed the snRNA-seq and snATAC-seq data. For each donor we removed duplicate reads from the ATAC bam files, converted to tagAlign format, and then quantified counts in 5kb windows tiling the human genome with bedtools^91^. We next cleaned background contamination from the RNA counts using SoupX^92^ (v1.6.2). We considered all barcodes with less than or equal to 500 unique RNA genes or less than or equal to 1000 unique ATAC fragments as empty droplets and all other barcodes as true cells to remove contamination from. We used the SoupX marker gene method for estimating contamination and used *INS*, *GCG*, *SST*, *PPY* as specific markers.

### Quality control of paired snRNA and snATAC-seq barcodes

To remove empty droplets, we remove barcodes with less than or equal to 500 unique RNA genes or less than or equal to 1000 unique ATAC fragments. We also removed data from all barcodes marked as multiplets, droplets that contained multiple gel beads, by 10x Genomics Cell Ranger ARC. After merging all samples together, further filters were applied to remove any remaining low-quality barcodes: we removed barcodes with more than 50,000 ATAC fragments, or more than 4,000 unique RNA genes, or greater than 1% of RNA mitochondrial gene counts, or more than 5,000 ATAC mitochondrial fragments, or a TSS enrichment score less than or equal to 2, as calculated by Seurat’s TSSEnrichment function.

### Doublet identification and removal methods

We used two different methods for identifying doublets. The first method was AMULET^93^, which uses snATAC-seq counts data to identify genomic loci with more than two haplotypes represented in a single barcode, indicating the barcode counts information comes from multiple cells. We converted the CellRanger Arc per_barcodes_metrics.csv file to one comparable with the CellRanger ATAC singlecell.csv output. Then we used the AMULET (v1.0) bash shell script with the ATAC bam file from CellRanger Arc and the --forcesorted option. The second method we used was Scrublet^94^, which simulates the profiles of potential doublets and then scores each barcode for their similarity to the simulated doublets. We ran Scrublet (v0.2.3) on the lightly filtered RNA counts data one sample at a time using min_counts=2, min_cells=3, min_gene_variability_pctl=85 and n_prin_comps=30. For all samples we used the minimum value between Scrublet’s automated doublet threshold score and 0.25 as a lower threshold to identify doublets. All barcodes from all samples identified as doublets by either AMULET or Scrublet were removed from all analyses.

### Clustering and cell type annotation of donor paired paired snRNA and snATAC-seq data

To merge counts data from all 28 Alberta samples, we combined post-SoupX normalized RNA counts and ATAC counts from 50k highly variable 5kb windows identified by applying the Seurat^26^ (v4.2) FindVariableFeatures command to all 5kb windows from 4 merged samples. To cluster the data, we first processed the RNA and ATAC data separately. For the RNA counts we used SCTransform^95^ to normalize the data, then performed PCA and UMAP^96^ with 50 dimensions to reduce the data dimensionality. For the ATAC counts we used term-frequency-inverse document frequency (TF-IDF)^97^ to normalize the count data, the FindTopFeatures command to identify the features with the most counts, and then singular value decomposition (SVD) and UMAP to reduce the data to lower dimensional space. We applied Harmony^98^ to the RNA and ATAC data separately to reduce batch effects, correcting across donor IDs. To combine the information from both the snRNA-seq and snATAC-seq assays, we used Seurat’s Weighted Nearest Neighbor algorithm^26^ and then reran UMAP and used the Leiden algorithm^99^ (resolution=0.25) to clusters the weighted nearest neighbor graph. After merging all donor samples into one object and clustering, we next called peaks with macs2^100^ using the ATAC counts from preliminary cell types defined from the windows clustering. Using a merged list of peaks from all cell types, we remade the ATAC counts matrix and then repeated our entire clustering procedure using peaks instead of 5kb windows. The cell type identity of all major clusters was manually annotated using known islet cell type marker genes.

### Peak calling on snATAC-seq data

We called peaks on ‘pseudo-bulk’ ATAC-seq profiles for each cell type defined from snATAC-seq data. In all cases these ‘pseudo’-bulk profiles reflect all barcodes assigned to a major cell type identified by clustering and the expression of marker genes. We combine the fragment information for all barcodes in the ‘pseudo’-bulk as tagAlign formatted files, and subsample these down to 100 million total fragments if necessary. We then use these as inputs for the macs2^100^ (v2.2.7.1) callpeak tool with an FDR cutoff of 0.05 and keeping all duplicates. Finally, we convert the default macs2 outputs to formats necessary for downstream analyses, including bigwigs. For downstream analyses, we limited the peak size to 300 bp and used an adjusted set of peaks, where overlapping peaks were represented by the peak with the highest accessibility across multiple cell types.

### Identifying cRE-gene links with Single-cell MultimOdal REgulatory Scorer (SMORES)

The SMORES method uses both snRNA-seq and snATAC-seq data to predict the target genes of cis-Regulatory Elements (cREs) in a cell type. SMORES takes a non-normalized RNA count matrix and a binarized ATAC count matrix as inputs, as well as a list of all ATAC peaks found in the cell type of interest and a list of genes expressed in the cell type of interest. For each gene that is expressed in the cell type, we perform the following steps: first, the gene expression per barcode is normalized by the sequencing depth of each barcode and scaled by 1M. Then all peaks within a 2Mbp window around the gene’s TSS are collected. For each peak-gene pair, non-parametric correlation is performed between the expression of the gene and the accessibility of the peak, using data from single cells. The barcode labels of both the gene expression and peak accessibility are shuffled randomly 100 times, and a peak-gene correlation value is calculated for each permutation. This creates a background distribution of correlation values that can be used to calculate an empirical p-value for the observed peak-gene correlation. After testing all gene-peak pairs, FDR correction is applied to the empirical p-values with Storey’s method and a paired bed file is output separately for all tested pairs and all significant pairs.

### Identifying cRE-gene links with ABC

We utilized the Activity-by-Contact^27^ (ABC) method to predict cRE-gene regulatory links. ABC is designed to use bulk ATAC-seq data as input, along with H3K27ac ChIP-seq and HiC data. To use it with our snATAC-seq data, we created ‘pseudo’-bulk profiles by merging the bam files for all barcodes assigned to a cell type. We then used CrossMap^101^ to liftover the ‘pseudo’-bulk bam files and cell type peak calls to hg19, to match the “BroadHiCdata” HiC reference data provided by ABC. Finally, we downloaded bulk islet and exocrine cell H3K27ac ChIP-seq peaks from Arda et al. (2018)^17^. We used the bulk adult endocrine data for all major islet cell types (beta, alpha, delta, and gamma cells) and combined adult and juvenile acinar and duct data for acinar and ductal cells, respectively. We ran ABC according to the recommended pipeline using these inputs and a list of genes expressed in each cell type (TPM>1), scaled the HiC data using the power law equation, and used the default threshold of 0.02. Finally, we used CrossMap to liftover ABC results to hg38 and processed them into paired bed files.

### Identifying cRE-gene links with Cicero

We used Cicero^28^ (v1.12.0) to predict cRE-gene regulatory links. For each cell type, we extracted a snATAC-seq counts matrix of all barcodes assigned to the cell type and all peaks called for the cell type. We then converted this matrix to a CellDataSet object and then a Cicero CDS object with aggregated information for k=30 nearby cells and ran Cicero with a window size of 1M bp, 100 max iterations, 100 sample regions, and a max distance constraint of 5M. To define significant Cicero links, we filtered for links that had a Cicero score greater than 0.05 and for which one of the cREs overlapped the TSS of at least one gene. Finally, we processed the resulting links into paired bed files.

### Comparing cRE-gene links found by SMORES, ABC, and Cicero

To evaluate the quality of cRE-gene links found by all methods, we compared their enrichment of two different measurements of islet regulatory activity: *cis*-expression quantitative trait loci (eQTLs) generated from bulk islet data by the InsPIRE consortium^16^ and 3D chromosome contacts from H3K27ac HiChIP data in primary human islets. For eQTLs, we first selected all links for which the cRE overlapped an eQTL and then calculated the enrichment of links with the same predicted target gene as the eQTL using a Fisher’s exact test. Next, for HiChIP we used the compare_connections function in Cicero to calculate how many cRE-gene links were between regions of interacting chromatin and then calculated the enrichment of overlapping links using a Fisher’s exact test. For comparisons between method specific distance-thresholded links, the background set was all links generated by the method within the distance bin that did not pass filtering thresholds. For comparisons between links found by multiple methods, the background set was all links generated by all three methods that did not pass filtering thresholds. Finally, we also calculated the enrichment of GWAS fine mapped variants impacting T2D risk^102^ and lead signals of variants impacting fasting glucose (FG) levels^9^ in cREs linked to target genes using finrich (v0.3.2). Briefly, this method empirically computes the enrichment of a set of variants in a list of cREs compared to a permuted background of all accessible peaks in the cell type.

### Identification of cell type-specific genes, cREs and TF motifs

We used Shannon entropy to define genes and cREs that had cell type specific expression and accessibility, respectively. Briefly, we first calculate normalized cell type ‘pseudo’-bulk measures of gene expression (TPM) and cRE accessibility (CPM). For cREs we used the set of 300bp peaks. We scaled the TPM and CPM values to probabilities between 0 and 1 by dividing each cell type measure by the sum of all cell type measures for that gene or cRE. We then calculated entropy for each gene and cRE using the Entropy() function from the DescTools package (v0.99.54)^103^. We classified all genes and peaks with an entropy value below 2 as cell type specific and assigned them to the cell type with the highest probability of expression or accessibility. Then, to identify motifs enriched in cell type specific cREs, we used HOMER’s^40^ findMotifsGenome.pl method on each set of cell type-specific cREs, using all cREs accessible in the cell type as a background set and a default region size of 200bp. Finally, to identify TFs likely binding these motifs in the cell type, we performed correlations between the expression of all family member TFs and the accessibility of the motif at the donor level. Family member TFs were derived from the TFClass database^104^, all correlations were non-parametric Spearman correlations, and two-sided p-values were computed using algorithm AS 89. We considered all cell type-specific TFs to be family member TFs which were significantly positively correlated (Benjamini Hochberg p-adjusted<.1) with a TF that was significantly enriched (Benjamini Hochberg p-adjusted<.1) in a cell type-specific cRE.

### Comparing sample cell type proportion to phenotypes

To compare the proportion of major cell types across samples with technical and donor covariates, we scaled the raw proportion value by taking the square root. We next trained a generalized linear model to predict these normalized counts per cell type by sample based on scaled versions of continuous per donor measurements: age, BMI and HbA1c, culture time, cold ischemia time, pancreas weight and purity. From this model we extracted the coefficient and p-value associated with each measurement. For the discrete measurements sex, collagenase type and pancreas consistency, we performed a Wilcoxon test for an association between each variable and the scaled proportion of cell types. We calculated a two-sided p-value for these results to determine if any associations were significant and corrected for multiple testing with the Benjamini Hochberg method. Donor R275 was excluded from these analyses as it was the only sample taken from a fibrotic pancreas. Donor R316 was excluded as it had lower islet purity (0.5) than other samples.

### Identification of genes and pathways changing with donor phenotypes

We used DESeq2^105^ (v1.34.0) to identify genes associated with specific phenotypes in both datasets separately. For each cell type, we created per-donor ‘pseudo’-bulk gene expression matrices, comprised of the sum of gene expression counts from all donor barcodes. Then we ran DESeq on each cell type using the data from all donors with more than 20 cells in the cell type and more than 1,000 total cells passing QC. DESeq was not run if less than 10 donors had enough cells. For DESeq we used two different formulas for the two datasets and all continuous variables were scaled prior to testing. When testing donors from Alberta, we corrected for all other per-donor phenotypes, culture time (hours), the proportion of beta cells in the donor and the mean number of genes expressed per barcode. When testing donors from HPAP, we corrected for all other per-donor phenotypes, the proportion of beta cells in the donor, tissue source, assay chemistry, self-identified race, and the mean number of genes expressed per barcode. HbA1c data was available for onle a subset of donors, so we only used those donors when performing DESeq2 for HbA1c and did not include HbA1c as a covariate in the formulas for other phenotypes.

Next, to extract trends occurring in both datasets, we performed a random effects-based meta-analysis using the metagen function from R’s meta package (v6.5-0)^106^. We used the “log2FoldChange” values from DESeq2 as effect size values and the “lfcSE” values as standard error, and the restricted maximum likelihood method to estimate between-study variance. After running meta-analysis separately on each gene, we performed FDR correction with Storey’s method and considered all genes with FDR<.1 as significantly associated to a phenotype in the cell type of interest.

We next used the FGSEA^107^ package (v1.20.0) to perform pre-ranked gene set enrichment analysis (GSEA) for each major cell type and phenotype using Gene Ontology, KEGG pathways, and Reactome terms. For these analyses we first removed ribosomal genes and then ranked genes up- and down-regulated using the gene’s -log10p-values multiplied by the effect size, both from meta-analysis. We considered terms as significant if they passed FDR<.1 and contained less than 500 genes.

### Processing of HPAP snATAC-seq data and cell type assignments

Single nuclear ATAC-seq fastq files for each non-diabetic HPAP donor were downloaded from the PANC-DB database^33,108^. 10x Genomics Cell ranger ATAC count (v2.0.0) was used to align reads to the GRCh38 reference genome from gencode v32. Then, we removed duplicate reads from bam files and generated read count matrices for 5kb windows tiling the genome for all 131,588 barcodes with a minimum of 1,000 ATAC reads. We next used this matrix to perform an initial clustering of the single cell data using Seurat^26^ (v4.2). Briefly, we first combined data from four samples, performed normalized with TF-IDF and then found the 50,000 most variable windows with Seurat’s FindTopFeatures method. We then used these 50,000 windows to combine data from all samples, performed TF-IDF and singular value decomposition, then we performed batch correction with Harmony^98^ and calculated a UMAP with the top 2-30 batch-corrected principal components. Finally, we identified clusters with the Leiden algorithm^99^ and 0.5 resolution. After clustering, we identified and removed 12,395 doublets using Amulet^93^, then reclustered the remaining barcodes and removed an additional 16,261 barcodes that had low FRiP (<= 0.3), low TSSe (<= 3), or were in clusters with overall low-quality metrics. We called peaks for each cluster using macs2^100^ and the same parameters described previously, and then recreated a peaks-based counts matrix for all barcodes. Then we removed 5,273 additional barcodes that were in small clusters with low-quality metrics and unclear marker genes. Our final snATAC-seq map contained 97,837 barcodes in total. Finally, cell types were annotated based on the expression of the same known islet marker genes used to annotate the multimodal data. For trait associations, per donor-‘pseudo’-bulk cell type matrices were generated using featureCounts^109^ and the same set of 300bp peaks called from our multimodal data.

### Identification of accessible chromatin regions changing with donor phenotypes

To identify accessible chromatin regions that change with phenotype, we implemented differential testing with DESeq on donors from both Alberta and HPAP together. For each cell type, we created per-donor ‘pseudo’-bulk accessible chromatin matrices, comprised of the sum of accessible chromatin counts from all donor barcodes for the set of fixed width peaks (300bp max). Then we ran DESeq on each cell type using the data from all donors with more than 20 cells in the cell type and more than 1,000 total cells passing QC. DESeq was not run if less than 10 donors had enough cells. For phenotype DESeq all continuous variables were scaled, and we corrected for all other per-donor phenotypes, culture time (hours), the proportion of beta cells in the donor and the average TSS enrichment (TSSe) per donor. We only had data on HbA1c from a subset of donors, so we only used those donors when performing DESeq2 for HbA1c and did not include HbA1c as a covariate in the models for other phenotypes.

Additionally, we used HOMER^40^’s findMotifsGenome script to identify transcription factor binding site (TFBS) motifs enriched in the same sets of differentially accessible peaks. For each HOMER analysis, we collected all results for known motifs and performed Benjamini Hochberg FDR correction. Finally, to test for concordance with the differential gene expression results for the same phenotypes, we performed fGSEA^107^ using pathways comprised of all genes predicted to be regulated by a phenotype-associated cRE (p<.01).

### Identification of TF motifs changing with donor phenotypes

To identify changes in TF motifs we ran chromVAR^39^ (v1.16.0) on the ATAC modality of our data. In brief, fixed width peak (300bp max) accessible chromatin count matrices were extracted and GRanges were constructed from the peak coordinates. Motif binding was predicted using motifmatchr^110^’s matchMotifs function (1.16.0) using BSgenome.Hsapiens.UCSC.hg38 as the reference and GC content was calculated via chromVAR’s addGCBias function. Following these steps the computeDeviations function was run. Single modality snATAC-seq data from the HPAP consortium was also run through chromVAR. A per barcode matrix matching the peaks used for chromVAR for the Alberta data was constructed and run through chromVAR using the same parameters as the Alberta data including the same motif match matrix.

To identify motifs with differential accessibility associated with age, sex, BMI, and HbA1c we ran linear mixed models on the chromVAR deviation scores matrix for each dataset independently using the lme4 (v1.1-35.2) and lmerTest (v3.1-3) packages. The formula motif ∼ HbA1c + scaled age + scaled BMI + sex + dataset + scaled culture time + scaled beta cell proportion + scaled TSS enrichment + scaled read depth in the ATAC modality + (1|samples) was used to test the effects of HbA1c and a model without HbA1c (due to lack of HbA1c information for some donors) for age, sex, and BMI. We performed a metanalysis of these models using the meta package (v7.0-0) to perform generic inverse variance meta-analysis with fixed effects.

We also employed HOMER^40^ to identify motifs enriched in cREs that change with phenotype. For each cell type and trait association, we took all cREs that were nominally significant (p<.01) from DESeq and split them based on the direction of change. Then for each set of cell type directionally changing cREs associated with a trait, we performed motif enrichment using the findMotifsGenome.pl from HOMER, with all peaks called in the cell type as background with the default region size (200bp) and re-parsing the hg38 genome file for this region size.

### Preparing Genotypes and Counts for caQTL Analysis

To prepare the imputed genotypes for caQTL analysis we filtered sample VCFs for SNPs based on imputation score (r^2^>0.7) using bcftools^111^ (v1.10.2). Sample genetic ancestry was identified by principal components analysis (PCA) using genotypes and ancestry from the 1000 genomes project^112^ as a reference.

For single cell caQTL identification and analyses, we followed a previously described approach based on ‘pseudo’-bulk quantifications^113^. Cell type assignments for each barcode were used to deconvoluted deduplicated sample .bam files into cell type specific, sample-specific .bam files using samtools^111^. The bam files were then used to make cell type ‘pseudo’-bulk count matrices using featurecounts^109^ (v2.0.0). A ‘pseudo’-bulk count matrix representing all peaks and barcodes was also generated to simulate bulk analysis. Count matrices were filtered for peaks containing an average of 5 reads per sample in the given cell type (or any cell type for bulk). Samples that were 3 or more standard deviations away from the mean in PCA analysis of the count matrices were removed from analysis for that cell type. Covariate matrices for each cell type were generated using the principle components identified by the make_covariates function in rasqualTools^41^ (v0.0.0.9000) as well as age and sex. Size factors were calculated and count matrices, covariate matrices and size factors were all converted to binary using rasqualTools.

Cell type and bulk specific VCFs were made by merging vcfs from the appropriate samples and filtering for SNPs within 100kb of a peak in the count matrix. Allele specific counts were added to these VCFs from bam files using creaseASVCF.sh from RASQUAL.

### caQTL Mapping

To perform caQTL analysis we tested for an association between each ATAC peak and SNPs within a 10kb cis-window using RASQUAL^41^, which performs joint QTL and allele specific effect (ASE) testing. We filtered out variants that are not well captured in our dataset using a minor allele frequency filter and feature SNP minor allele frequency filter of 0.05. An empirical FDR of 5% was calculated using q-values from two permutation tests to adjust q-value FDR cutoffs. Lead SNPs for each peak were identified as the lowest p-value SNP tested, with the SNP closest to the center of the peak being chosen in the case of a tie in p-value. While caQTLs were mapped for all major cell types, endothelial, immune, and stellate cells were excluded from subsequent analyses due to the limited power associated with the rarity of these cell types in our dataset and the lack of significant associations detected.

### Comparison to Bulk Islet caQTLs

Due to the heterogeneous nature of islets, we compared ‘pseudo’-bulk and cell type results to identify cell type specific effects. We used the package mashr (v0.2.79)^42^ to estimate the cell type specificity of caQTLs, using a lfsr cutoff of 0.05. We also annotated cRE cell type specificity based on a CPM filter of >6. We also identified cREs with significant caQTL associations in each cell type that were not identified in our bulk like analysis.

Additionally, we compared our cell type caQTLs to a dataset of caQTLs derived from bulk ATAC-seq analysis of 19 islet samples^14^. First, we mapped feature locations of the significant caQTL summary statistics from Khetan et al. from hg19 to hg38 using liftOver and identified overlapping features (ATAC peaks, called OCRs in by Khetan et al.) using bedtools^91^ intersect. SNPs were mapped from hg19 to hg38 by rsID using data from dbGap release 156. From this intersection we identified feature SNP pairs tested in both analyses for comparison. Percent concordance was calculated as the percent of features sharing the same direction of effect (both <0.5 or both >0.5). We also performed this analysis comparing cell type specific effects without bulk-like analysis.

### caQTL Transcription Factor Motif Analysis

To examine transcription factor binding motifs disrupted by caQTL SNPs, we subset the set of lead caQTL SNPS to only those found within the feature of interest, as it is less clear how a motif outside of the feature of interest may be altering chromatin accessibility. These SNPs were tested against motifs of human transcription factors in JASPAR2022 from MotifDb^114^ (v1.36.0) using motifbreakR^43^ (v2.8.0) using the “ic” method. Enrichment of disrupted motifs in each cell type was measured using motifbreakR (v2.16) to identify JASPAR motifs from motifDB (v1.44) disrupted by any caQTL lead variant (lowest p value) within the caQTL feature regardless of significance. We performed a Fisher’s test on each motif in each cell type to identify motifs whose predicted disruption was enriched in variants with a significant caQTL association. We also identified motifs disrupted by significant caQTL lead variants outside of any peaks. We used bedtools intersect to identify significant lead variants outside of any peak in our dataset. Then we used motifbreakR to identify distrusted motifs for all lead caQTL variants and used a Fisher’s test to identify disrupted motifs enriched in variants outside of peaks.

### Genetic datasets

We used T1D summary statistics and credible sets previously published by our group^13^ as the basis for T1D risk. For T2D, we downloaded summary statistics and the multi-ancestry credible set from the DIAMANTE consortium^115^. For all glycemic traits (glycated hemoglobin: HbA1c, fasting insulin: FI, fasting glucose: FG, 2-h glucose after an oral glucose challenge: 2hGlu) we downloaded summary statistics from the MAGIC consortium and lead variants from their 2021 study^9^. To construct quasi-credible sets for each glycemic trait, we used their lead variants for every locus and the identified all SNPs with high LD to these variants (R^2^ > 0.8) using the LDproxy tool from the LDlinkR package (v1.4.0)^116,117^. For each locus, we then calculated an equally weighted PIP for all variants within high LD.

### Enrichment of GWAS signal in genomic regions

To calculate the enrichment of GWAS signal in sets of genomic regions we used the FINRICH method^76^. Briefly, FINRICH measures the overlap of variants in a credible set with a set of genomic regions of interest, and then measures the significance of this overlap by comparing to the overlap of the same credible set with a permuted background set of equal size. Each variant overlap was weighted by the PIP of the variant from fine mapping and 1000 permutations were used to calculate an empirical p value for the overlap. For enrichments of GWAS signal in trait associated cREs, we used a background set of all peaks that were tested in trait associations. For testing the enrichment of GWAS signal in caQTL cREs, we used all cREs that were called peaks by MACS2 as a background set.

### Colocalization of caQTLs with GWAS

First, we identified caQTLs where the lead SNP was within 100 kb of a lead SNP passing genome-wide significance in GWAS for type 1 diabetes^13^, type 2 diabetes^115^, or the glycemic traits glycosylated hemoglobin (HbA1c), fasting glucose (FG), 2-h glucose after an oral glucose challenge (2hGlu) and fasting insulin (FI)^9^. For these caQTLs, we re-ran caQTL analysis with the matrixEQTL tool^118^ (v2.3) to give sufficient genomic coverage for statistical colocalization using a window of 100kb, quantile normalized counts, no minor allele frequency filter, and the same covariates as RASQUAL. The caQTL association results were then colocalized with the summary statistics from each GWAS study using the coloc (v5.2.3) method^119^. Briefly, we prepared datasets for each trait and caQTL feature which included the following information for each SNP and both datasets: effect (beta), variance of effect, p values, minor allele frequency, and number of individuals in the study that generated the dataset. Then we performed colocalization with the approximate Bayes Factor method and assumed that there is only one causal variant per trait. We initiated the model with the default priors for potential outcomes: only the GWAS has a genetic association (1e-4), only the caQTL has a genetic association (1e-4), and both traits are associated and share a single causal variant (1e-5). All loci with the probability of a shared causal variant greater than 0.8 were considered as colocalized.

## Author contributions

H.M. and W.E. performed single cell and genetic analyses and wrote the manuscript. K.K., R.E., and E.G. contributed to single cell analyses. P.B. and A.J. contributed to genetic analyses. P.K. performed single cell analyses and created tools and visualizations for single cell data. M.M. performed single cell assays. J.E.M.F. contributed to donor sample processing and preparation. M.I.M. contributed to genetic analyses and obtaining funding. S.P. contributed to the generation of single cell assays. A.L.G. contributed to genetic analyses and obtaining funding. P.E.M. contributed islet donor samples and phenotypic information and to obtaining funding. K.J.G. supervised the study, obtained funding, contributed analyses, and wrote the manuscript.

## Data availability

Raw data will be deposited in GEO upon publication. Single cell maps and associated resources such as gene regulatory programs and associations are available at http://multiome.isletgenomics.org. All other data and results are provided with the manuscript or available from the authors on request.

## Code availability

All code used in our study can be found in the GitHub repository https://github.com/Gaulton-Lab/non-diabetic-islet-multiomics.

## Acknowledgements

Human islets for research were obtained with the assistance of the Human Organ Procurement and Exchange (HOPE) program, Trillium Gift of Life Network (TGLN), and other Canadian organ procurement organizations. Islet isolation was approved by the Human Research Ethics Board at the University of Alberta (Pro00013094). All donor families gave informed consent for the use of pancreatic tissue in research. Data was also obtained from the Human Pancreas Analysis Program (HPAP-RRID:SCR_016202), a Human Islet Research Network (RRID:SCR_014393) consortium (UC4-DK-112217, U01-DK-123594, UC4-DK-112232, and U01-DK-123716). The work in this study was supported by NIH grants DK105554, DK138512, and HG012059 and an Endowed Chair in Type 1 Diabetes Research to K.J.G. Work in Alberta was supported in part by a grant from the Canadian Institutes of Health Research to P.E.M. (PS 186226). P.E.M. holds the Canada Research Chair in Islet Biology. H.M.M. was supported by the DT O’Connor Scholar in Genetics and W.E. was supported by the T32 GM008666 training grant.

## Conflicts of interest

K.J.G. has done consulting for Genentech, received honoraria from Pfizer, and is a shareholder of Neurocrine biosciences. M.I.M. is currently an employee of Genentech, and a holder of Roche stock.

## Notes

http://multiome.isletgenomics.org

