## Supplemental figures for "Single cell multiome profiling of pancreatic islets reveals physiological changes in cell type-specific regulation associated with diabetes risk"

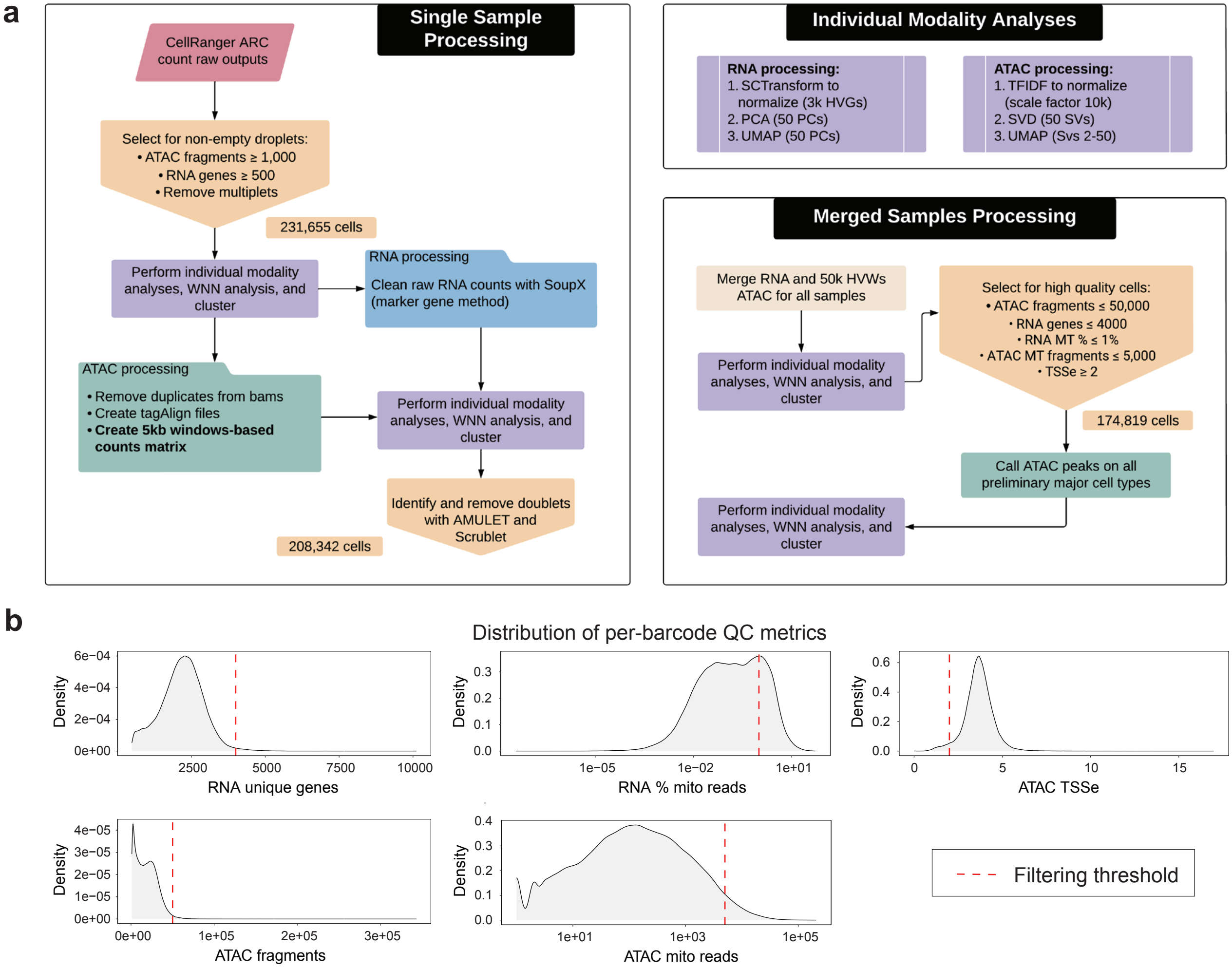


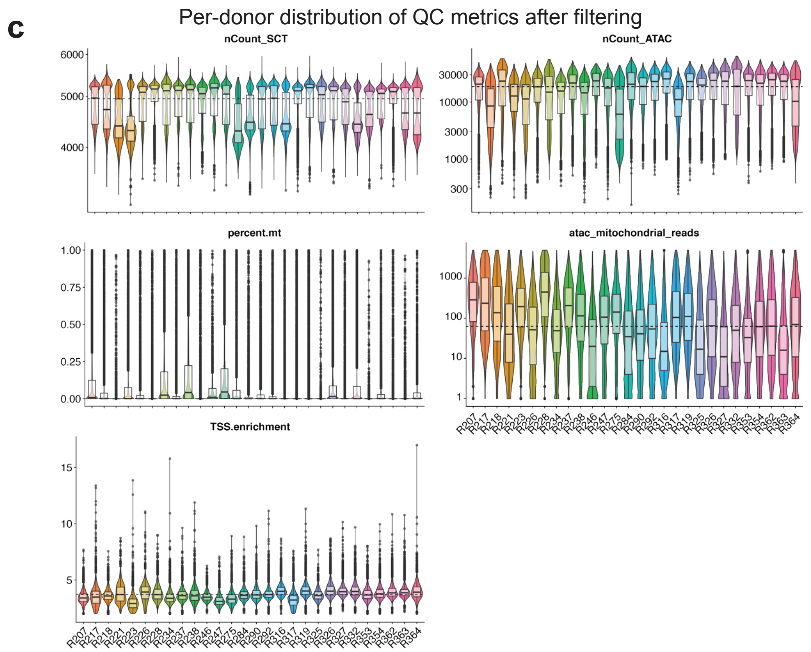


**d**

| Donor | Amulet doublets | Scrublet doublets | All doublets | Final # cells | Proportion beta cells | SoupX contamination |
|---|---|---|---|---|---|---|
| R207 | 679 | 347 | 867 | 6248 | 0.52 | 0.088 |
| R217 | 1006 | 667 | 1489 | 5619 | 0.25 | 0.038 |
| R218 | 507 | 164 | 619 | 2172 | 0.69 | 0.1474 |
| R221 | 367 | 237 | 515 | 4902 | 0.22 | 0.0265 |
| R223 | 706 | 572 | 1150 | 6343 | 0.54 | 0.0867 |
| R226 | 416 | 371 | 621 | 5869 | 0.55 | 0.0733 |
| R228 | 772 | 385 | 1014 | 4041 | 0.23 | 0.0512 |
| R234 | 394 | 509 | 713 | 7458 | 0.41 | 0.0483 |
| R237 | 413 | 343 | 586 | 6211 | 0.44 | 0.0315 |
| R238 | 496 | 317 | 720 | 6186 | 0.69 | 0.1363 |
| R246 | 165 | 97 | 210 | 1878 | 0.30 | 0.0222 |
| R247 | 561 | 336 | 743 | 5341 | 0.40 | 0.037 |
| R275 | 940 | 142 | 1045 | 3430 | 0.51 | 0.1947 |
| R284 | 503 | 486 | 792 | 6082 | 0.24 | 0.0237 |
| R290 | 564 | 636 | 922 | 8009 | 0.37 | 0.0422 |
| R292 | 510 | 343 | 709 | 6749 | 0.54 | 0.0697 |
| R316 | 963 | 928 | 1464 | 10752 | 0.45 | 0.0357 |
| R317 | 495 | 385 | 780 | 8531 | 0.61 | 0.1325 |
| R319 | 709 | 481 | 972 | 8942 | 0.67 | 0.083 |
| R325 | 149 | 107 | 203 | 1823 | 0.42 | 0.0481 |
| R326 | 808 | 869 | 1247 | 9798 | 0.49 | 0.0308 |
| R327 | 509 | 321 | 686 | 6820 | 0.63 | 0.0445 |
| R332 | 1125 | 615 | 1477 | 7488 | 0.23 | 0.0344 |
| R353 | 380 | 470 | 646 | 7025 | 0.46 | 0.0598 |
| R354 | 470 | 337 | 649 | 5945 | 0.40 | 0.0349 |
| R362 | 529 | 311 | 667 | 7529 | 0.67 | 0.1106 |
| R363 | 696 | 751 | 1038 | 9469 | 0.42 | 0.0418 |
| R364 | 644 | 168 | 772 | 4159 | 0.33 | 0.0269 |

**Supplementary Figure 1. Multimodal single cell data processing and quality control metrics.** a) Overview flowchart of our single sample and merged sample processing pipeline for 10x Multiome data. b) Density plots representing the spread of QC metrics in all non-empty droplets, after AMULET doublet removal. The dotted red lines mark the metric cutoffs chosen to remove low quality data barcodes. For RNA unique genes, RNA % mito reads, ATAC fragments and ATAC mito reads these were upper cutoffs; for ATAC TSSe this was a minimum cutoff. c) Violin plots of per-donor distribution of QC metrics after final QC filters were applied. d) Donor level summary of doublets removed, number of barcodes included in the final map, proportion of beta cells and SoupX estimated contamination fraction.

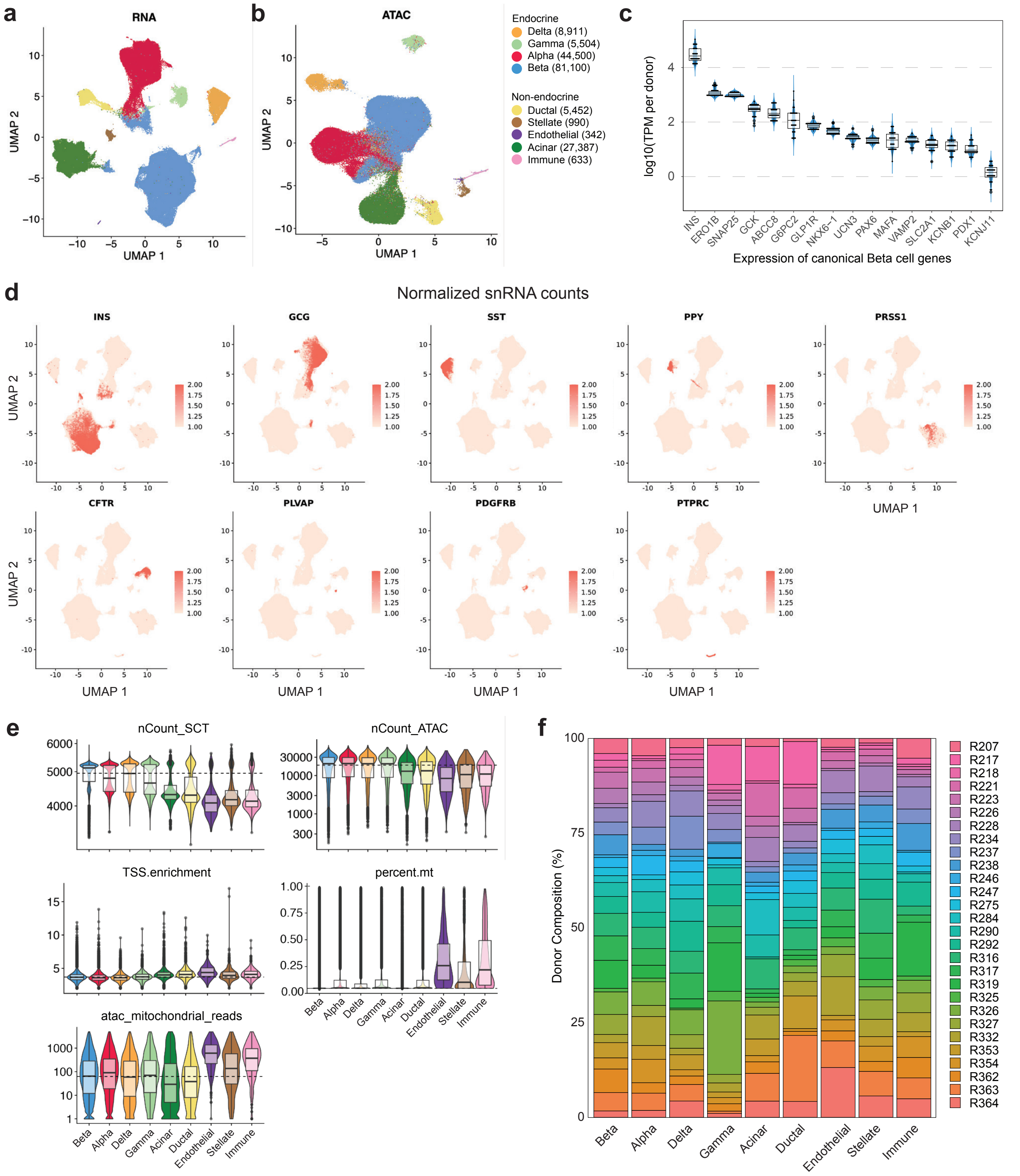
a
RNA
UMAP 1
UMAP 2
b
ATAC
UMAP 1
UMAP 2
Endocrine
Delta (8,911)
Gamma (5,504)
Alpha (44,500)
Beta (81,100)
Non-endocrine
Ductal (5,452)
Stellate (990)
Endothelial (342)
Acinar (27,387)
Immune (633)
c
log10(TPM per donor)
INS
ERO1B
SNAP25
GCK
ABCC8
G6PC2
GLP1R
NKX6-1
UCN3
PAX6
MAFA
VAMP2
SLC2A1
KCNB1
PDX1
KCNJ11
Expression of canonical Beta cell genes
d
Normalized snRNA counts
INS
GCG
SST
PPY
PRSS1
CFTR
PLVAP
PDGFRB
PTPRC
UMAP 1
UMAP 2
e
nCount_SCT
nCount_ATAC
TSS.enrichment
percent.mt
atac_mitochondrial_reads
Beta
Alpha
Delta
Gamma
Acinar
Ductal
Endothelial
Stellate
Immune
f
Donor Composition (%)
R207
R217
R218
R221
R223
R226
R228
R234
R237
R238
R246
R247
R275
R284
R290
R292
R316
R317
R319
R325
R326
R327
R332
R353
R354
R362
R363
R364

**Supplementary Figure 2. Single modality clustering and characterizations of cell type assignments.** a-b) Single modality UMAPs of single nuclear RNA (a) and ATAC (b) based clustering of all final barcodes. c) Range of per-donor ‘pseudo’-bulk beta cell TPM values for an extended list of canonical beta cell markers. d) SCT-normalized expression of cell type marker genes projected onto the combined modality UMAP. e) Per-cell type distribution of QC metrics after final QC filters were applied. f) Percentage of barcodes from each donor for all cell types.

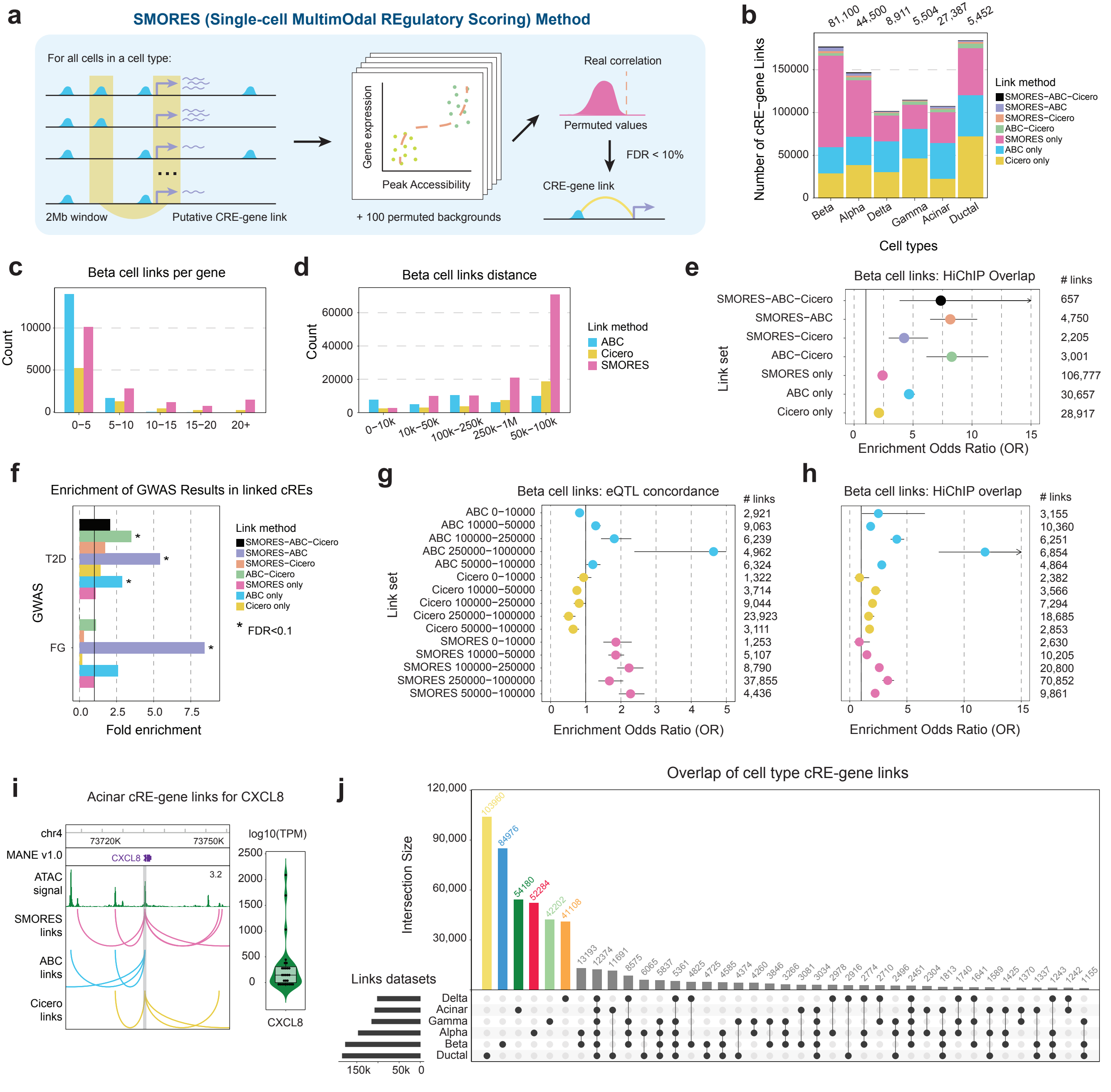
a
SMORES (Single-cell MultimOdal REgulatory Scoring) Method
For all cells in a cell type:
2Mb window
Putative CRE-gene link
Gene expression
Peak Accessibility
+ 100 permuted backgrounds
Real correlation
Permuted values
FDR < 10%
CRE-gene link
b
Number of cRE-gene Links
Cell types
Beta
Alpha
Delta
Gamma
Acinar
Ductal
81,100
44,500
8,911
5,504
27,387
5,452
Link method
SMORES−ABC−Cicero
SMORES−ABC
SMORES−Cicero
ABC−Cicero
SMORES only
ABC only
Cicero only
c
Beta cell links per gene
Count
0−5
5−10
10−15
15−20
20+
d
Beta cell links distance
Count
0−10k
10k−50k
100k−250k
250k−1M
50k−100k
Link method
ABC
Cicero
SMORES
e
Beta cell links: HiChIP Overlap
Link set
Enrichment Odds Ratio (OR)
### links
SMORES−ABC−Cicero 657
SMORES−ABC 4,750
SMORES−Cicero 2,205
ABC−Cicero 3,001
SMORES only 106,777
ABC only 30,657
Cicero only 28,917
f
Enrichment of GWAS Results in linked cREs
GWAS
T2D
FG
Fold enrichment
Link method
SMORES−ABC−Cicero
SMORES−ABC
SMORES−Cicero
ABC−Cicero
SMORES only
ABC only
Cicero only
* FDR<0.1
g
Beta cell links: eQTL concordance
Link set
Enrichment Odds Ratio (OR)
### links
ABC 0−10000 2,921
ABC 10000−50000 9,063
ABC 100000−250000 6,239
ABC 250000−1000000 4,962
ABC 50000−100000 6,324
Cicero 0−10000 1,322
Cicero 10000−50000 3,714
Cicero 100000−250000 9,044
Cicero 250000−1000000 23,923
Cicero 50000−100000 3,111
SMORES 0−10000 1,253
SMORES 10000−50000 5,107
SMORES 100000−250000 8,790
SMORES 250000−1000000 37,855
SMORES 50000−100000 4,436
h
Beta cell links: HiChIP overlap
Enrichment Odds Ratio (OR)
### links
3,155
10,360
6,251
6,854
4,864
2,382
3,566
7,294
18,685
2,853
2,630
10,205
20,800
70,852
9,861
i
Acinar cRE-gene links for CXCL8
chr4
73720K
73750K
MANE v1.0
CXCL8
ATAC signal
3.2
SMORES links
ABC links
Cicero links
log10(TPM)
CXCL8
j
Overlap of cell type cRE-gene links
Intersection Size
Links datasets
Delta
Acinar
Gamma
Alpha
Beta
Ductal
103960
84976
54180
52284
42202
41108
13193
12374
11691
8575
6065
5837
5361
4825
4725
4585
4374
4260
3846
3266
3081
3034
2978
2916
2774
2710
2496
2451
2304
1813
1740
1641
1589
1425
1370
1337
1243
1242
1155

**Supplementary Figure 3. Evaluation of prediction methods for cRE-gene target links.** a) Overview of Single-cell MultimOdal Regulatory Scoring Method (SMORES), for each cREs within a 2Mb window centered around the gene of interest the correlation between binarized cRE accessibility and the gene is calculated and then compared to the correlation generated by 100 permuted barcode sets to generate an empirical p-value. b) Summary of the number of cRE-gene links predicted by each method, or a combination of methods, for each cell type with more than 1,000 barcodes. c-d) Distribution of beta cell cRE-gene links predicted by each method based on number of links per gene (c) and the distance between the cRE and gene TSS (d). e) Acinar cell cRE-gene links between CXCL8 and all cREs predicted to regulate it, separated by prediction method. Right inlay shows the range of per-donor 'pseudo'-bulk acinar cell TPM values for CXCL8. f) Enrichment of eQTLs with the same target gene prediction in cREs of sets of links separated by method and distance between cRE and gene TSS. g-h) Enrichment of overlapping HiChIP genomic contacts in sets of links separated by method and distance (g) and separated by prediction methods (g). i) Enrichment of GWAS credible sets link sets of cRE-gene links separated by prediction methods, as calculated by FINRICH. Stars indicate enrichments passing FDR<.1. j) Overlap of all cell type cRE-gene links sets between cell types, for each cell type we are comparing all links generated by any method. The upset plot shows the 40 different cell type overlap combinations with the most links.

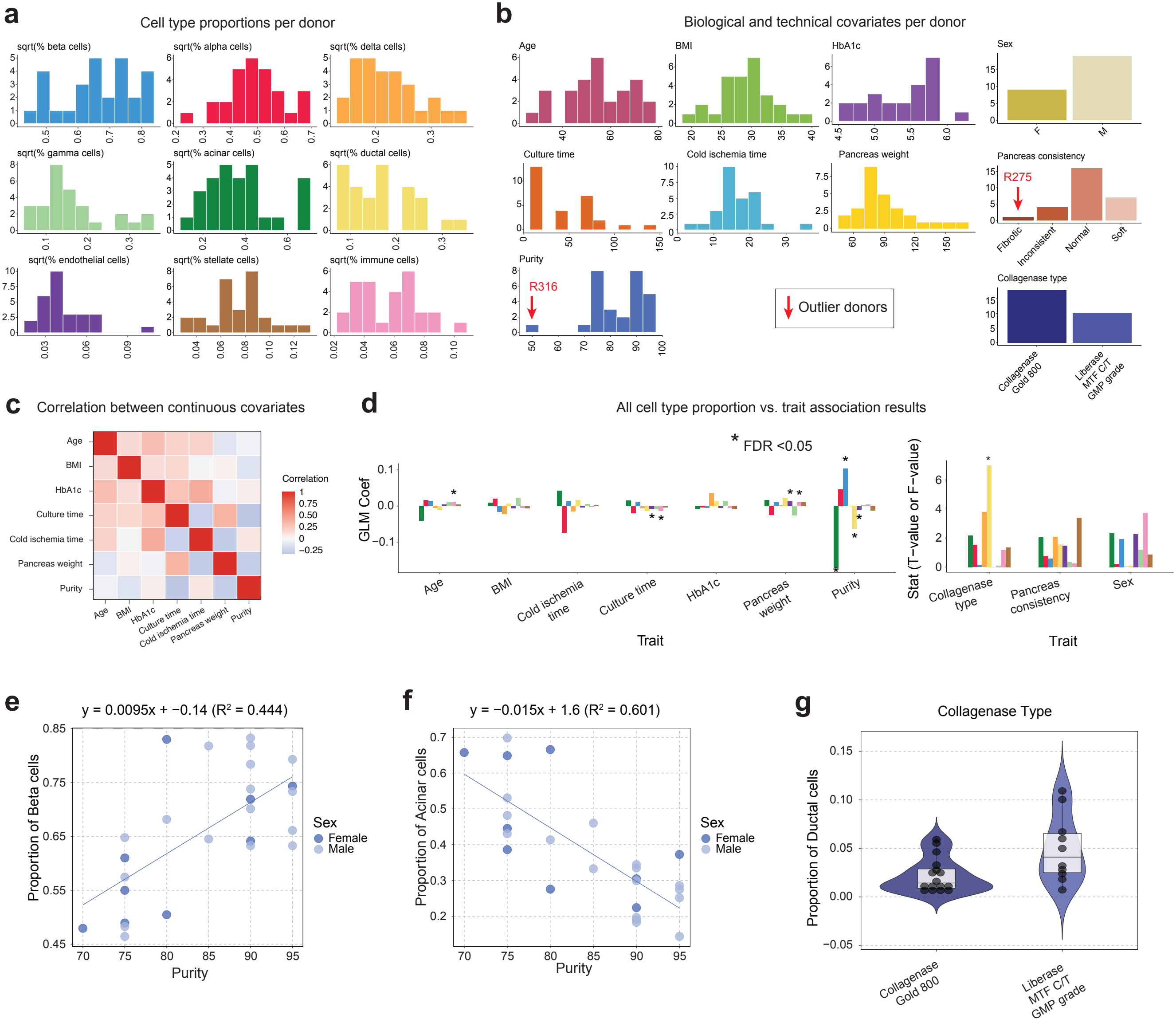

a
Cell type proportions per donor
sqrt(% beta cells)
sqrt(% alpha cells)
sqrt(% delta cells)
sqrt(% gamma cells)
sqrt(% acinar cells)
sqrt(% ductal cells)
sqrt(% endothelial cells)
sqrt(% stellate cells)
sqrt(% immune cells)
b
Biological and technical covariates per donor
Age
BMI
HbA1c
Sex
Culture time
Cold ischemia time
Pancreas weight
Pancreas consistency
R275
Fibrotic
Inconsistent
Normal
Soft
Purity
R316
Outlier donors
Collagenase type
Collagenase Gold 800
Liberase MTF C/T GMP grade
c
Correlation between continuous covariates
Correlation
d
All cell type proportion vs. trait association results
* FDR <0.05
GLM Coef
Stat (T−value or F−value)
Trait
e
y = 0.0095x + −0.14 (R² = 0.444)
Proportion of Beta cells
Purity
Sex
Female
Male
f
y = −0.015x + 1.6 (R² = 0.601)
Proportion of Acinar cells
g
Collagenase Type
Proportion of Ductal cells

**Supplementary Figure 4. Comparisons between cell type proportions and donor covariates.** a) Distribution of the proportion of each cell type across donors. Proportion values were scaled by taking the square root of the raw proportion. b) Distribution of biological and technical covariates across all Alberta donors, outlier donors that were not included in proportion associations are indicated with a red arrow. c) Heatmap of the correlation between all continuous covariates. d) Summary of all cell type proportion versus biological and technical covariates associations. For associations with continuous covariates, the coefficient of effect from the linear model is plotted, and for associations with discrete covariates the F value from the ANOVA test is plotted. Significant associates (FDR<.05) are marked with *. e-f) The proportion of beta (e) and acinar (f) cells compared to islet isolation purity. Donor points are colored based on sex: female (dark blue), male (light blue) and a basic linear equation was fit to the relationship, with the equation displayed on top of the graph. g) Distribution of the proportion of ductal cells amongst samples treated with two different collagenase types.

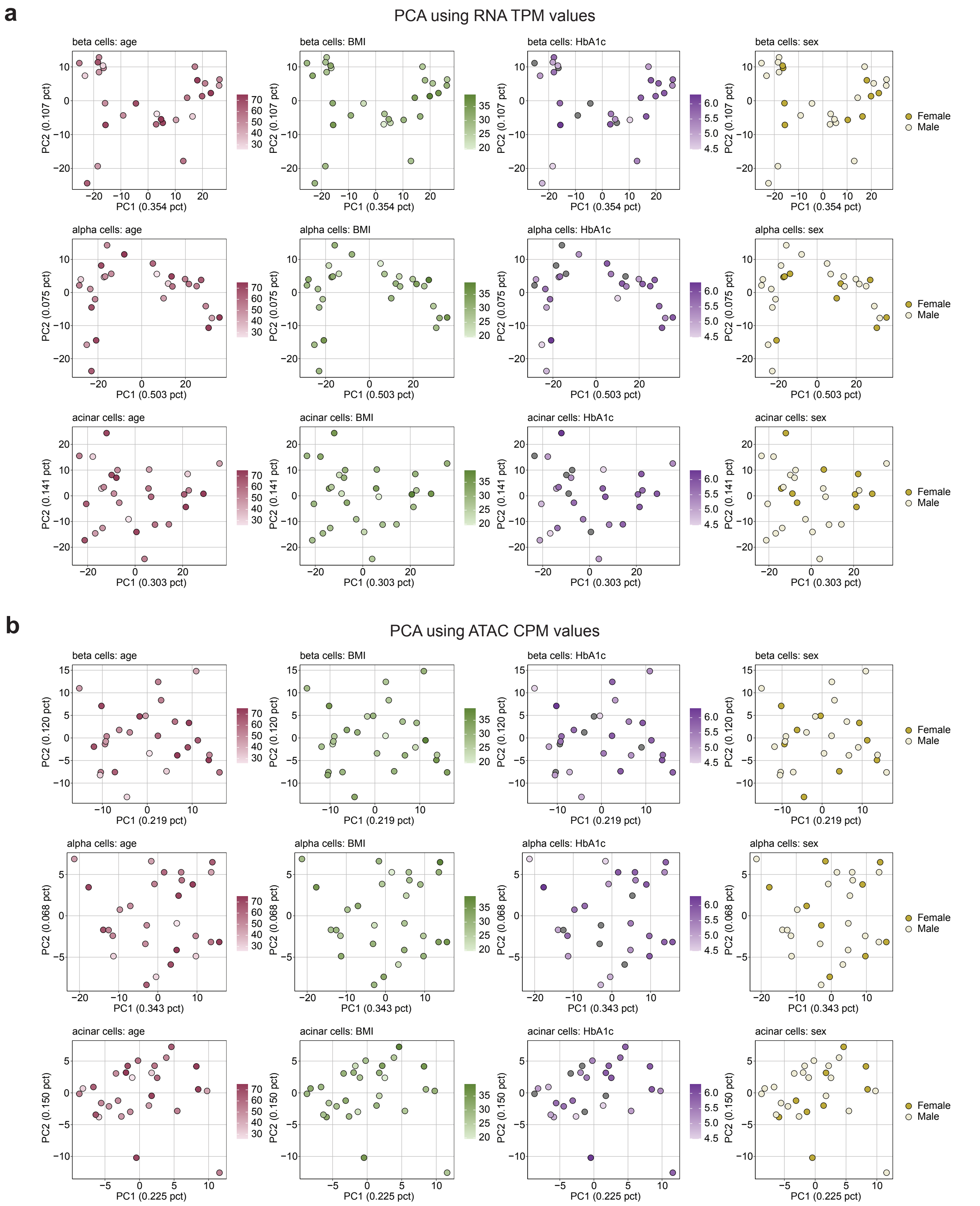
a
PCA using RNA TPM values
beta cells: age
beta cells: BMI
beta cells: HbA1c
beta cells: sex
PC1 (0.354 pct)
PC2 (0.107 pct)
alpha cells: age
alpha cells: BMI
alpha cells: HbA1c
alpha cells: sex
PC1 (0.503 pct)
PC2 (0.075 pct)
acinar cells: age
acinar cells: BMI
acinar cells: HbA1c
acinar cells: sex
PC1 (0.303 pct)
PC2 (0.141 pct)
Female
Male
b
PCA using ATAC CPM values
beta cells: age
beta cells: BMI
beta cells: HbA1c
beta cells: sex
PC1 (0.219 pct)
PC2 (0.120 pct)
alpha cells: age
alpha cells: BMI
alpha cells: HbA1c
alpha cells: sex
PC1 (0.343 pct)
PC2 (0.068 pct)
acinar cells: age
acinar cells: BMI
acinar cells: HbA1c
acinar cells: sex
PC1 (0.225 pct)
PC2 (0.150 pct)
Female
Male

**Supplementary Figure 5. Principal component analysis of donor gene expression and chromatin accessibility.** a) Donors plotted based on the first and second principal components after performing PCA on per-donor beta, alpha and acinar cell gene expression TPM values. The plots in each column are colored by a different phenotype. b) Donors plotted based on the first and second principal components after performing PCA on per-donor beta, alpha and acinar cell chromatin accessibility CPM values. The plots in each column are colored by a different phenotype.

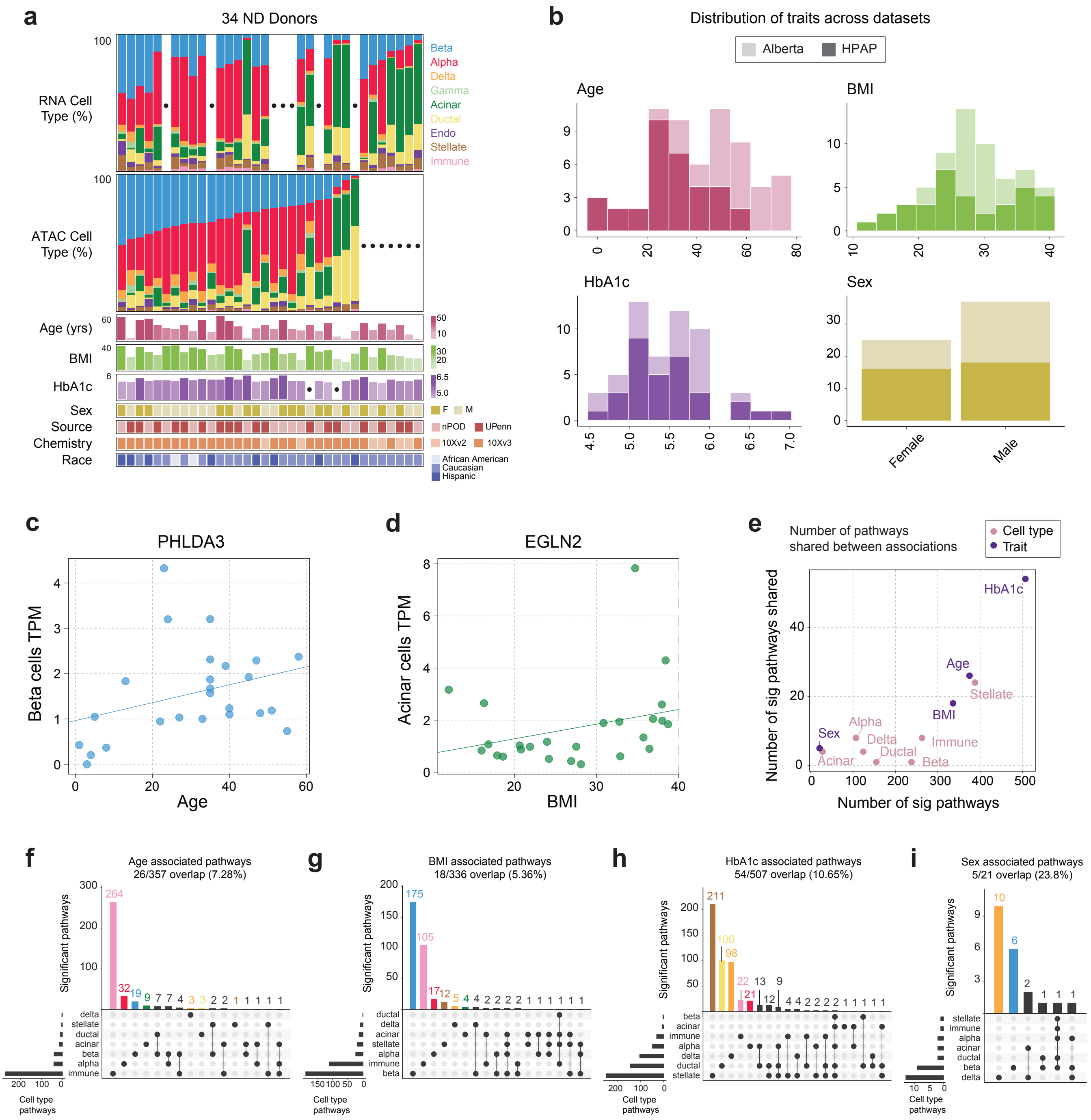
a
34 ND Donors
RNA Cell Type (%)
ATAC Cell Type (%)
Beta
Alpha
Delta
Gamma
Acinar
Ductal
Endo
Stellate
Immune
Age (yrs)
BMI
HbA1c
Sex
Source
Chemistry
Race
F
M
nPOD
UPenn
10Xv2
10Xv3
African American
Caucasian
Hispanic
b
Distribution of traits across datasets
Alberta
HPAP
Age
BMI
HbA1c
Sex
Female
Male
c
PHLDA3
Beta cells TPM
Age
d
EGLN2
Acinar cells TPM
BMI
e
Number of pathways shared between associations
Cell type
Trait
Number of sig pathways shared
Number of sig pathways
HbA1c
Age
Stellate
BMI
Sex
Alpha
Delta
Ductal
Immune
Acinar
Beta
f
Age associated pathways
26/357 overlap (7.28%)
Significant pathways
Cell type pathways
g
BMI associated pathways
18/336 overlap (5.36%)
h
HbA1c associated pathways
54/507 overlap (10.65%)
i
Sex associated pathways
5/21 overlap (23.8%)

**Supplementary Figure 6. Summary of phenotypes in the HPAP dataset and gene expression associations.** a) Distribution of cell type proportions and phenotype information across all 34 non-diabetic HPAP donors, where each column represents a donor. The dots indicate donors with missing measurements, where 17 of the donors had both RNA and ATAC measurements. b) Distribution of phenotypes in donors compared between the two datasets, Alberta (light colors) and HPAP (dark colors). c-d) 'Pseudo'-bulk per-donor TPM values for PHLDA3 (c) and EGLN2 (d) for donors from the HPAP dataset, plotted against age (c) and BMI (d). e) Summary of the overlap in all pathways associated with a trait or a cell type. The numbers of pathways refer to the total number of unique pathways associated with a trait across cell types, or a cell type across traits. Points are labeled based on whether the comparison is of all pathways per a trait (purple), or per a cell type (pink). f-i) Upset plots illustrating the overlap of different pathways associated between cell types and age (f), BMI (g), HbA1c (h) and sex (i).

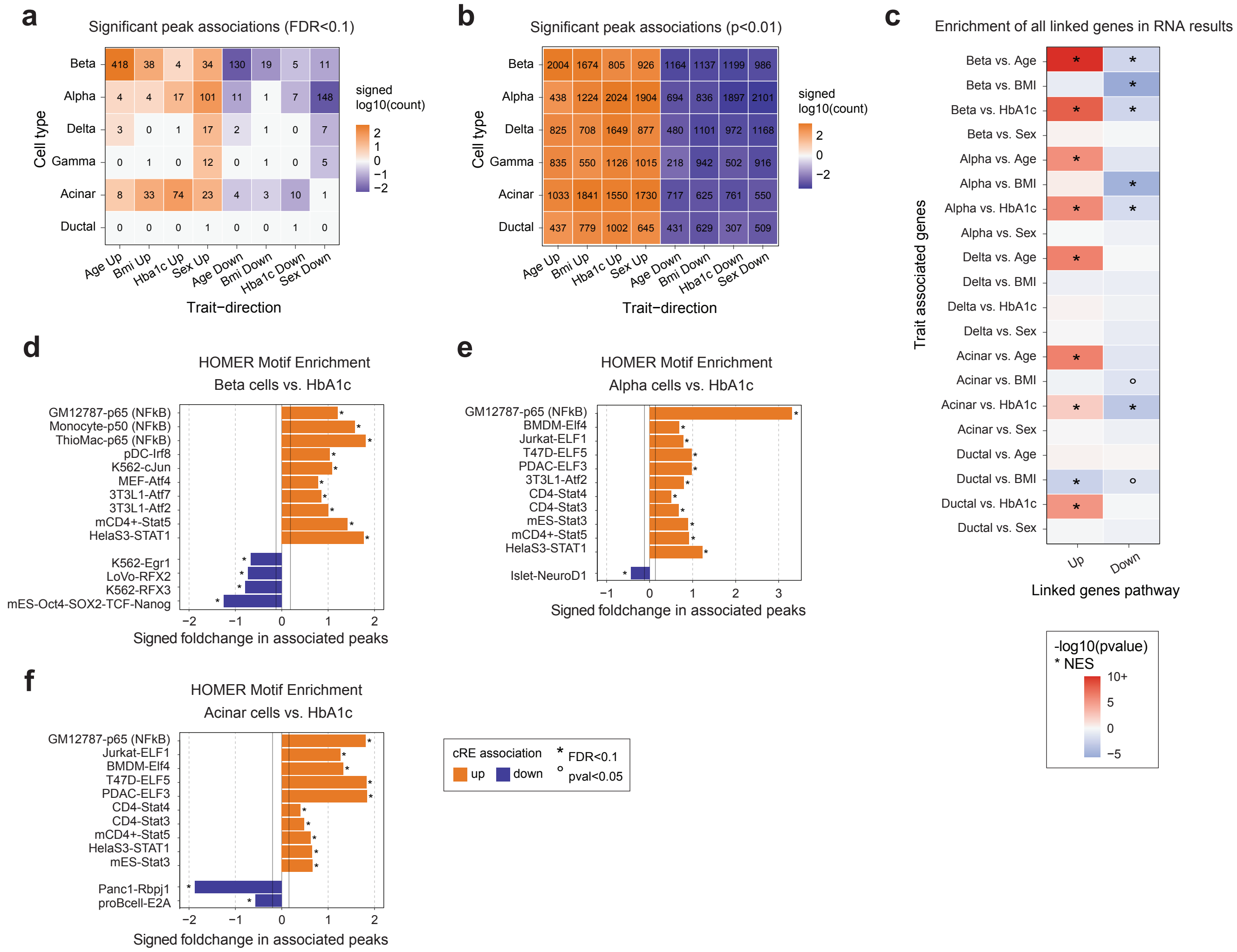
a
Significant peak associations (FDR<0.1)
Beta
Alpha
Delta
Gamma
Acinar
Ductal
Cell type
signed log10(count)
Age Up
Bmi Up
Hba1c Up
Sex Up
Age Down
Bmi Down
Hba1c Down
Sex Down
Trait−direction
b
Significant peak associations (p<0.01)
c
Enrichment of all linked genes in RNA results
Trait associated genes
Linked genes pathway
Up
Down
-log10(pvalue)
* NES
d
HOMER Motif Enrichment
Beta cells vs. HbA1c
Signed foldchange in associated peaks
e
HOMER Motif Enrichment
Alpha cells vs. HbA1c
f
HOMER Motif Enrichment
Acinar cells vs. HbA1c
cRE association
up
down
* FDR<0.1
° pval<0.05

**Supplementary Figure 7. Changes in chromatin accessibility associated with donor phenotypes.** a-b) Summary of the number of significantly differentially accessible (a; FDR <.1) and nominally differentially accessible (b; p<.01) cREs in associations between cell types and traits. Groups of cREs were divided based on the direction of differential accessibility and then the total number of differential cREs per association was scaled with log10 and signed based on the direction of change. c) Enrichment of all target genes of cREs nominally associated with a phenotype for a cell type in differentially expressed genes associated with the corresponding phenotype. Enrichments were performed by using fGSEA with a database of pathways comprised of all genes linked to a phenotype-associated cRE, split by direction of association. The color of each block is based on the product of the -log10 p-value and the normalized enrichment score (NES) from fGSEA. Nominal (p<.05, °) and FDR significant (FDR<.1,*) enrichments are marked. d-f) HOMER motif enrichment results for cREs increasing (orange) or decreasing (purple) with HbA1c in beta (d), alpha (e) or acinar (f) cells. The solid lines denote the foldchange value associated with the top 20% of motifs tested. Nominal (p<.05, °) and FDR significant (FDR<.1,*) enrichments are marked.

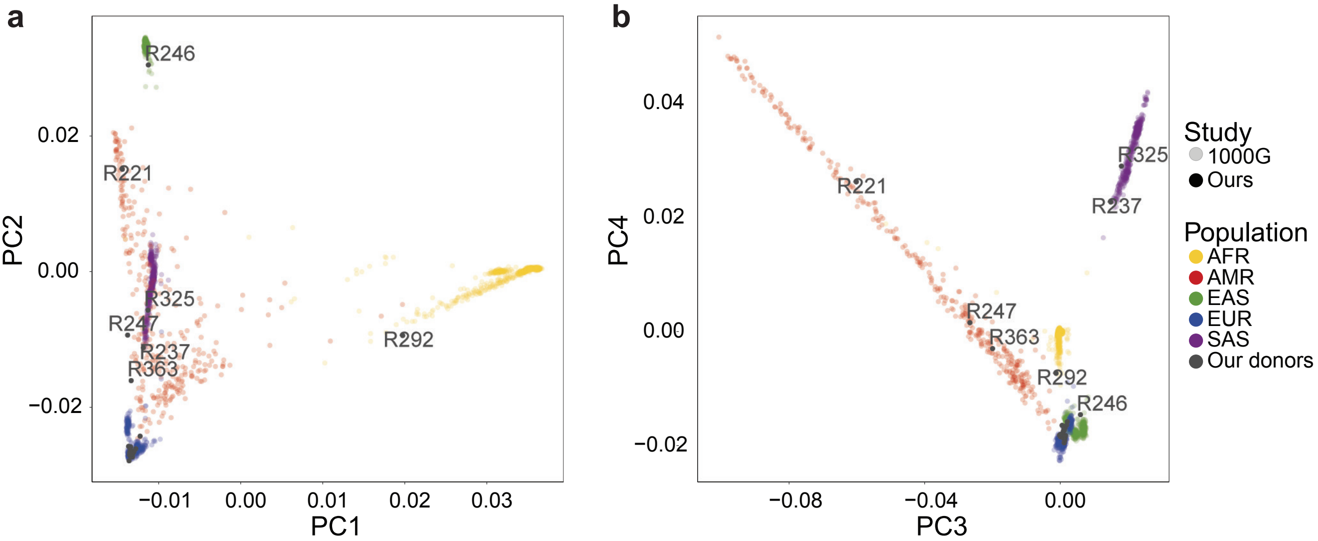


**c** Summary of caQTL results

| Cell type | caQTL FDR 10% | caQTL FDR 5% | caQTL FDR 1% | Tested features |
|---|---|---|---|---|
| **Bulk** | 16034 | 12217 | 7455 | 252057 |
| **Beta** | 7789 | 5932 | 3657 | 106506 |
| **Alpha** | 5629 | 4218 | 2546 | 107935 |
| **Delta** | 1262 | 877 | 478 | 79636 |
| **Gamma** | 758 | 524 | 274 | 60138 |
| **Acinar** | 3050 | 2144 | 1011 | 96373 |
| **Ductal** | 419 | 267 | 106 | 49135 |
| **Endothelial** | 0 | 0 | 0 | 1399 |
| **Stellate** | 2 | 2 | 2 | 11915 |
| **Immune** | 9 | 4 | 4 | 8173 |

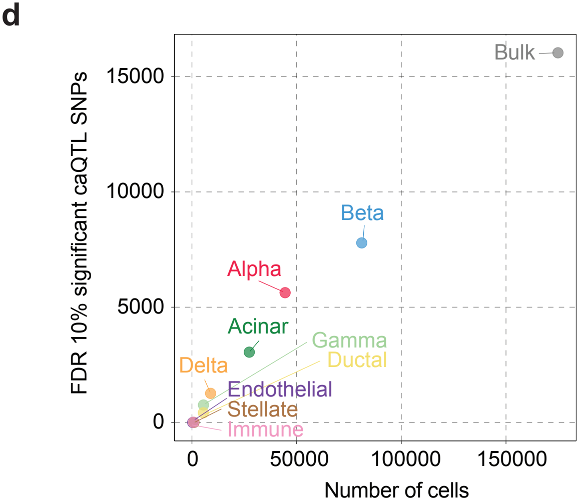

**Supplementary Figure 8. Donor ancestries and summary of caQTL results.** a-b) Genotype PCA plots for PCs 1 and 2 (a) and PCs 3 and.4 (b). Reference genotypes from the 1000 Genomes project[102] are included and colored according to ancestry group. All donors from our study are colored grey and all non-European donors are label with donor ID. c) Summary of caQTLs passing different FDR thresholds, and how many features were initially tested per cell type. d) Comparison between the number of cells in the cell type and the number of significant caQTL SNPs (FDR<.05) found in the cell type.

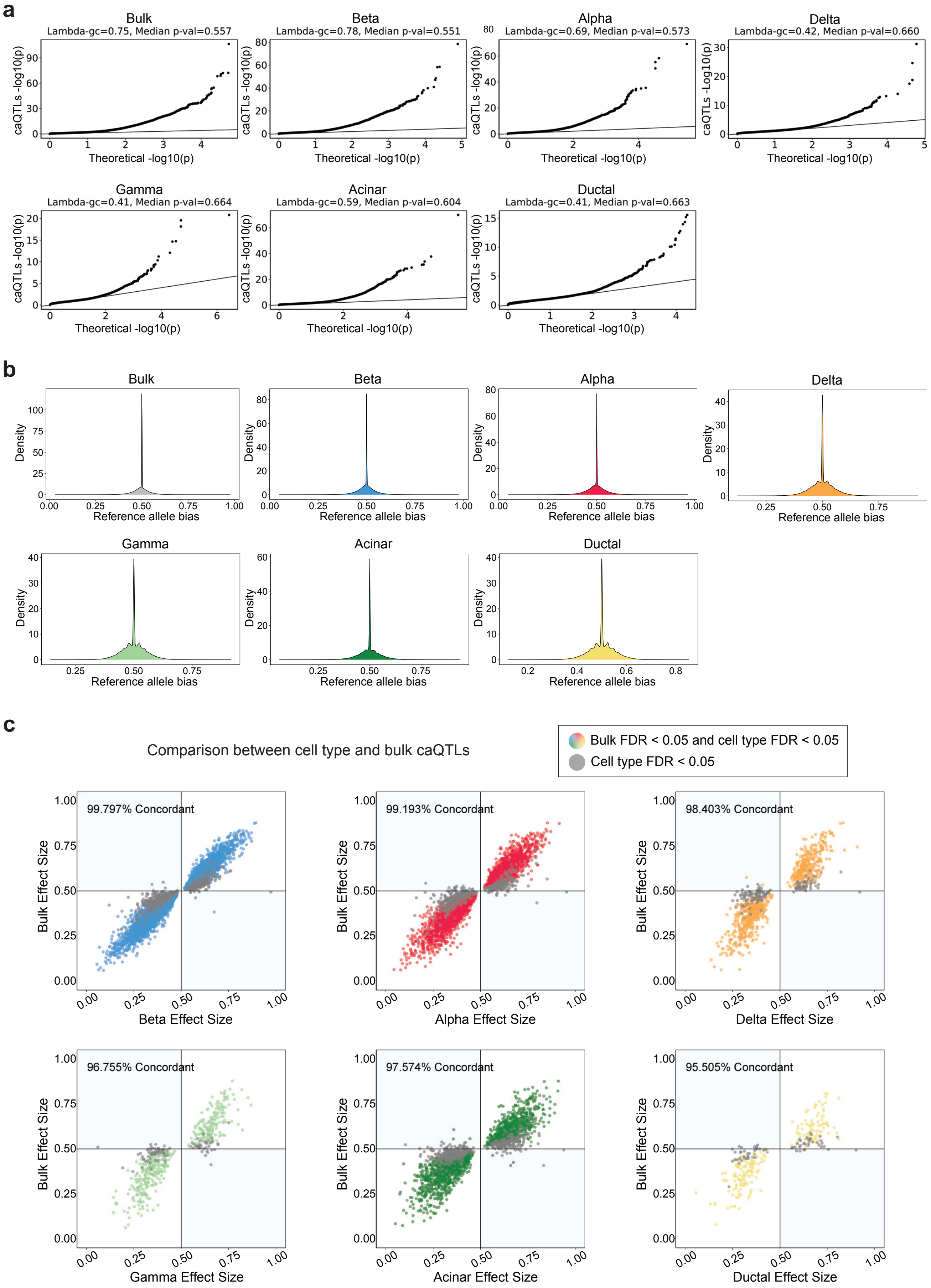

a
Bulk
Lambda-gc=0.75, Median p-val=0.557
Beta
Lambda-gc=0.78, Median p-val=0.551
Alpha
Lambda-gc=0.69, Median p-val=0.573
Delta
Lambda-gc=0.42, Median p-val=0.660
Gamma
Lambda-gc=0.41, Median p-val=0.664
Acinar
Lambda-gc=0.59, Median p-val=0.604
Ductal
Lambda-gc=0.41, Median p-val=0.663
caQTLs -log10(p)
Theoretical -log10(p)
b
Bulk
Beta
Alpha
Delta
Gamma
Acinar
Ductal
Density
Reference allele bias
c
Comparison between cell type and bulk caQTLs
Bulk FDR < 0.05 and cell type FDR < 0.05
Cell type FDR < 0.05
99.797% Concordant
99.193% Concordant
98.403% Concordant
96.755% Concordant
97.574% Concordant
95.505% Concordant
Bulk Effect Size
Beta Effect Size
Alpha Effect Size
Delta Effect Size
Gamma Effect Size
Acinar Effect Size
Ductal Effect Size

**Supplementary Figure 9. caQTL quality control metrics and comparison of cell type caQTLs to bulk.** a) Per cell type qq-plots for all caQTL p-values compared to a theoretical uniform distribution. The inflation factor, lambda-GC, and the median p-value for each cell type are listed above each plot. b) Density plots of the distribution of reference allele bias in the caQTLs for all cell types. c) Comparison of caQTL effect size (x-axis) and 'pseudo'-bulk islet caQTL effect sizes (y-axis). All SNPs that are significant cell type caQTLs are included and are colored based on whether they pass FDR significance in the 'pseudo'-bulk. The percentage of variants with concordant directional effects is listed at the top of each plot.

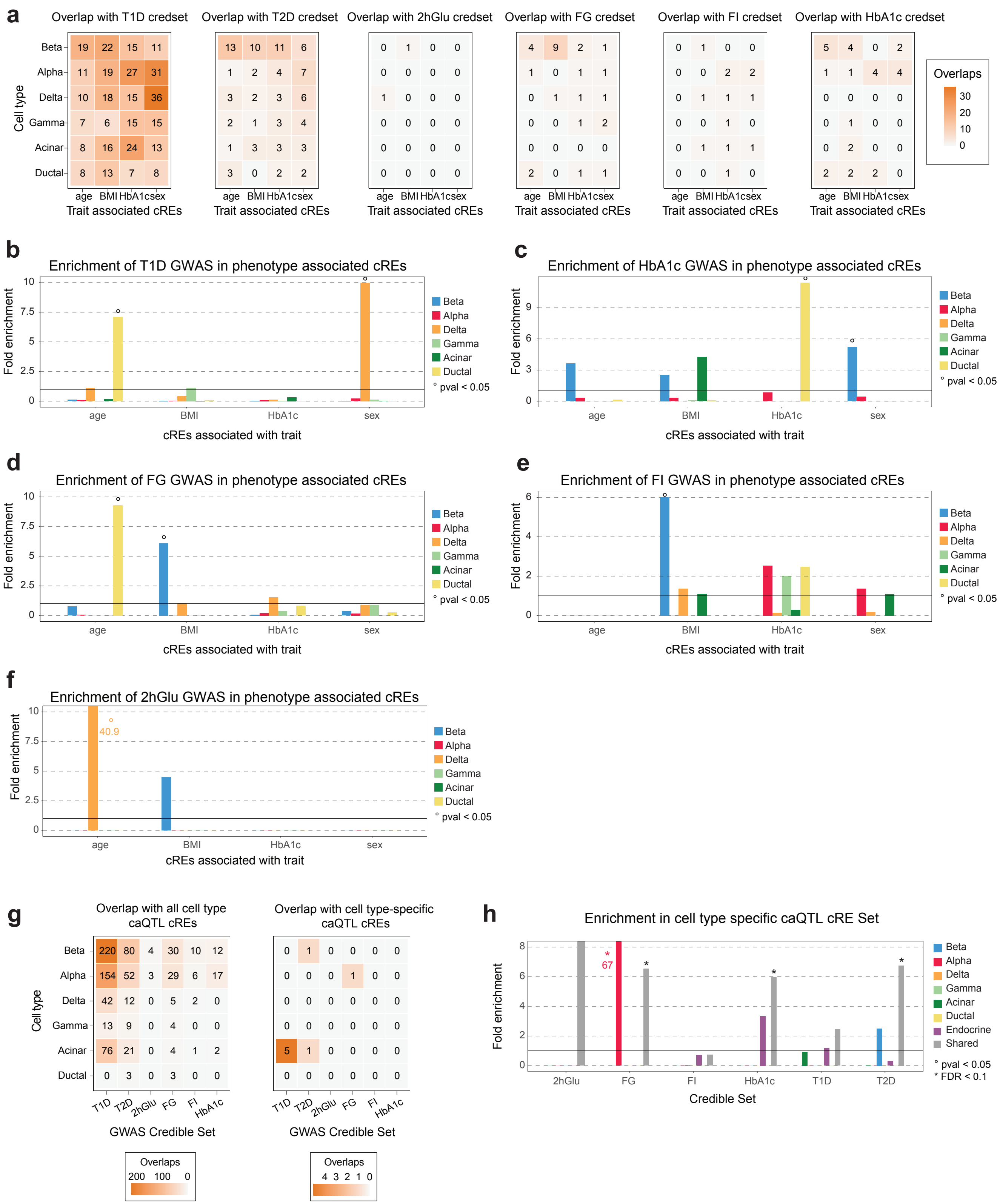

a
Overlap with T1D credset
Overlap with T2D credset
Overlap with 2hGlu credset
Overlap with FG credset
Overlap with FI credset
Overlap with HbA1c credset
Cell type
Beta
Alpha
Delta
Gamma
Acinar
Ductal
age BMI HbA1c sex
Trait associated cREs
Overlaps
b
Enrichment of T1D GWAS in phenotype associated cREs
c
Enrichment of HbA1c GWAS in phenotype associated cREs
d
Enrichment of FG GWAS in phenotype associated cREs
e
Enrichment of FI GWAS in phenotype associated cREs
f
Enrichment of 2hGlu GWAS in phenotype associated cREs
Fold enrichment
cREs associated with trait
° pval < 0.05
40.9
g
Overlap with all cell type caQTL cREs
Overlap with cell type-specific caQTL cREs
GWAS Credible Set
h
Enrichment in cell type specific caQTL cRE Set
Endocrine
Shared
Credible Set
* FDR < 0.1
67

**Supplementary Figure 10. Overlap of GWAS credible sets and cREs associated with phenotypes and genotypes.** a) Summary of the number of cREs associated with traits that overlap the T1D, T2D, and glycemic trait credible sets. b-f) Enrichment of SNPs in the T1D (b), HbA1c (c), fasting glucose (d), fasting insulin (e), and 2-h glucose after an oral glucose challenge (f) credible sets that overlap cREs associated with age, BMI, HbA1c and sex, in the six largest cell types. Nominally significant enrichments (pval<.05) are indicated with °. g) Summary of the number of cREs with one or more significant caQTLs that overlap GWAS credible sets. h) Enrichment of SNPs in the glycemic trait, T1D and T2D credible set that overlap cREs that have a cell type-specific, lineage-specific, or shared significant caQTL in each cell type. FDR significant enrichments (FDR<.1) are indicated with * and nominally significant enrichments (pval<.05) are indicated with °.

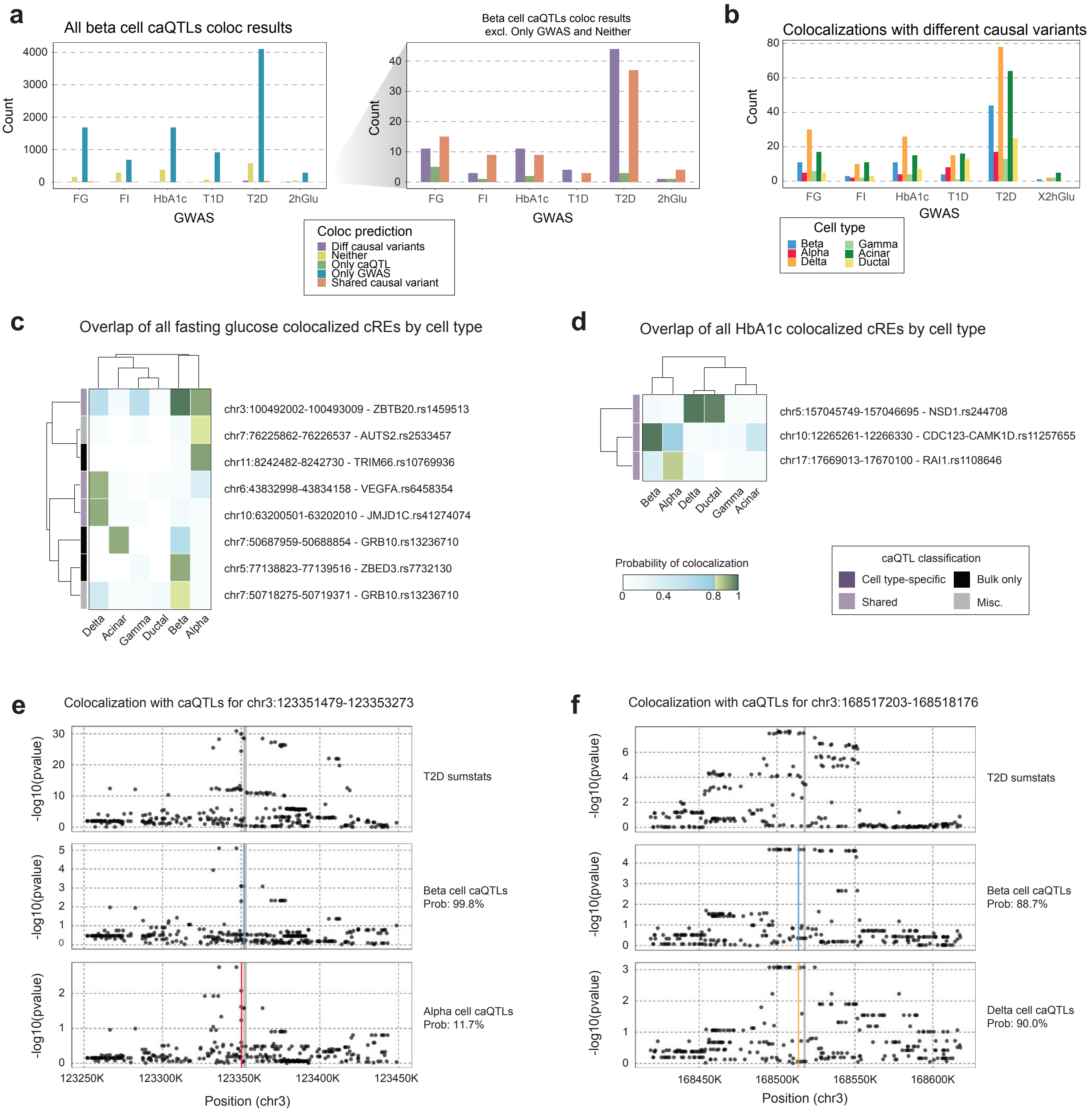

a
All beta cell caQTLs coloc results
Count
GWAS
FG
FI
HbA1c
T1D
T2D
2hGlu
Beta cell caQTLs coloc results
excl. Only GWAS and Neither
Coloc prediction
Diff causal variants
Neither
Only caQTL
Only GWAS
Shared causal variant
b
Colocalizations with different causal variants
X2hGlu
Cell type
Beta
Alpha
Delta
Gamma
Acinar
Ductal
c
Overlap of all fasting glucose colocalized cREs by cell type
chr3:100492002-100493009 - ZBTB20.rs1459513
chr7:76225862-76226537 - AUTS2.rs2533457
chr11:8242482-8242730 - TRIM66.rs10769936
chr6:43832998-43834158 - VEGFA.rs6458354
chr10:63200501-63202010 - JMJD1C.rs41274074
chr7:50687959-50688854 - GRB10.rs13236710
chr5:77138823-77139516 - ZBED3.rs7732130
chr7:50718275-50719371 - GRB10.rs13236710
Delta
Acinar
Gamma
Ductal
Beta
Alpha
d
Overlap of all HbA1c colocalized cREs by cell type
chr5:157045749-157046695 - NSD1.rs244708
chr10:12265261-12266330 - CDC123-CAMK1D.rs11257655
chr17:17669013-17670100 - RAI1.rs1108646
Probability of colocalization
caQTL classification
Cell type-specific
Bulk only
Shared
Misc.
e
Colocalization with caQTLs for chr3:123351479-123353273
-log10(pvalue)
T2D sumstats
Beta cell caQTLs
Prob: 99.8%
Alpha cell caQTLs
Prob: 11.7%
123250K
123300K
123350K
123400K
123450K
Position (chr3)
f
Colocalization with caQTLs for chr3:168517203-168518176
T2D sumstats
Beta cell caQTLs
Prob: 88.7%
Delta cell caQTLs
Prob: 90.0%
168450K
168500K
168550K
168600K
Position (chr3)

**Supplementary Figure 11. Statistical colocalization between caQTLs and GWAS credible sets.** a) Summary of all statistical colocalization results between beta cell caQTLs and all credible sets. For each beta cell cRE, the colocalization prediction with the highest PIP was assigned as the prediction, and what the colored bars represent. Right: zoomed in plot for just colocalizations of different causal variants, only the caQTL has a causal variant, or shared causal variants. b) Summary of the number of statistically significant (PIP>0.8) loci with evidence for different causal variants between cell type caQTLs and summary statistics for glycemic traits, T1D, and T2D. c-d) The PIP for each fasting glucose (c) and HbA1c (d) colocalized cRE across all cell types. Non-significant colocalizations are colored in blue and significant colocalizations are colored in green. Row names indicate the nearest credible set to the cRE that overlaps GWAS SNPs. e-f) Examples of loci where one or two cell-type caQTLs (e:chr3-123,351,479-123,353,273; f:chr3:168,517,203-168,518,176) colocalized with T2D risk. The lead SNP for each cell type caQTL is indicated by the colored line.

**Supplementary Tables and Data**

**Supplementary Table 1. Characteristics of donors profiled in this study**
**Supplementary Table 2. Expression level of genes in pancreatic cell types**
**Supplementary Table 3. Cis-regulatory elements in pancreatic cell types**
**Supplementary Table 4. Genes with cell type-specific specific expression levels**
**Supplementary Table 5. Cis-regulatory elements with cell type-specific activity**
**Supplementary Table 6. Sequence motifs enriched in cell type-specific cREs**
**Supplementary Table 7. Significant correlations of sample variables with principal components**
**Supplementary Table 8. Donors from HPAP consortium used in study**
**Supplementary Table 9. Associations between cell type proportions and donor covariates**
**Supplementary Table 10. Genes associated with phenotypes in pancreatic cell types**
**Supplementary Table 11. Gene set enrichment of phenotype-associated genes**
**Supplementary Table 12. Cis-regulatory elements associated with phenotypes in pancreatic cell types**
**Supplementary Table 13. Sequence motif accessibility associated with phenotypes in pancreatic cell types**
**Supplementary Table 14. Sequence motifs enriched in cREs associated with phenotypes**
**Supplementary Table 15. Sequence motifs enriched in caQTLs for pancreatic cell types**
**Supplementary Table 16. Enrichment of GWAS signals in sets of phenotype-associated cREs**
**Supplementary Table 17. Enrichment of GWAS signals in sets of genotype-associated cREs**
**Supplementary Table 18. Diabetes and glycemic trait loci with pancreatic cell type caQTLs**
**Supplementary Table 19. Diabetes and glycemic trait loci colocalized with pancreatic cell type caQTLs**

**Supplementary Data 1. Links between cis-regulatory elements and target genes**
**Supplementary Data 2. Summary statistics of chromatin QTLs in pancreatic cell types**
**Supplementary Data 3. Re-estimated chromatin QTL effects in pancreatic cell types**
